# PD-L1 activity is associated with partial EMT and metabolic reprogramming in carcinomas

**DOI:** 10.1101/2022.10.05.510941

**Authors:** Srinath Muralidharan, Manas Sehgal, R Soundharya, Susmita Mandal, Sauma Suvra Majumdar, M Yeshwanth, Aryamaan Saha, Mohit Kumar Jolly

**Affiliations:** Centre for BioSystems Science and Engineering, Indian Institute of Science, Bangalore 560012, India; Department of Biotechnology, National Institute of Technology, Durgapur 713216, India; Department of Biotechnology, Indian Institute of Technology Madras, Chennai 600036, India

**Keywords:** Partial EMT, Meta-analysis, Immune checkpoint molecules, Glycolysis, Oxidative phosphorylation, Metabolic plasticity

## Abstract

Immune evasion and metabolic reprogramming are hallmarks of cancer progression often associated with a poor prognosis and frequently present significant challenge for cancer therapies. Recent studies have emphasized on the dynamic interaction between immunosuppression and the dysregulation of energy metabolism in modulating the tumor microenvironment to promote cancer aggressiveness. However, a pan-cancer association among these two hallmarks, and a potent common driver for them – Epithelial-Mesenchymal Transition (EMT) – remains to be done. Here, our meta-analysis across 184 publicly available transcriptomic datasets as well as The Cancer Genome Atlas (TCGA) data reveals that an enhanced PD-L1 activity signature along with other immune checkpoint markers correlate positively with a partial EMT and elevated glycolysis signature but a reduced OXPHOS signature in many carcinomas. These trends were also recapitulated in single-cell RNA-seq time-course EMT induction data across cell lines. Furthermore, across multiple cancer types, concurrent enrichment of glycolysis and PD-L1 results in worse outcomes in terms of overall survival as compared to enrichment for only PD-L1 activity or expression. Our results highlight potential functional synergy among these interconnected axes of cellular plasticity in enabling metastasis and/or multi-drug resistance in cancer.

## Introduction

The dynamic interplay between host immunity and cancer cells often facilitates tumor development and is now recognized as a cancer hallmark [1,2]. Cancer cells exploit a myriad of counter-regulatory mechanisms which are crucial for self-tolerance, to limit the host immune attack, creating a tumor-promoting immune-suppressive milieu [3,4]. Many immune checkpoint molecules such as PD-L1 [5], CTLA4 [6], CD276 [7], LAG3 [8] and HAVCR2 [9] are known to block anti-tumor responses in the tumor microenvironment [10]. Particularly, the programmed death receptor-ligand 1 (PD-L1) plays a crucial role in the metastatic spread of tumors by binding to programmed death receptor 1 (PD-1) on CD8+ T cells, thus imparting potent immune evasive character to cancer cells mainly by altering effector functions of T cells, along with inhibition of T cell proliferation and survival [11,12]. Accumulating evidence suggests that cancer cells can exhibit high PD-L1 expression [13–15] and this upregulation can be influenced by multiple signaling pathways [16,17]. Thus, an understanding of diverse array of regulatory mechanisms that govern immune evasion and their implications for therapeutic intervention in cancer is an active area of research.

The upregulation of PD-L1-mediated immunosuppression by cancer cells has been extensively associated with epithelial-mesenchymal transition (EMT), a process often implicated in metastasis [18–22]. Furthermore, cells with a hybrid epithelial/mesenchymal (E/M) phenotype can also express elevated levels of PD-L1 [19,23]. On a different note, emerging evidence supports the notion that in pathological conditions such as cancer, cells undergoing EMT also exhibit varying degrees of metabolic reprogramming [23]. Alterations in key metabolic pathways such as glycolysis, oxidative phosphorylation, lipid metabolism and amino acid metabolism influence cancer progression at least partly by modulating the EMT status of cells [25,26]. More recently, metabolic reprogramming has been increasingly reported to be inter-linked with immune evasion [1,27,28]. For instance, increased glycolytic metabolism of cancer cells favors cancer growth by competing with T cells for growth and/or blocking lactic acid export in T cells, thus impeding cytotoxic activity of T cells [28,29]. Concomitantly, inhibition of PD-L1 expression when coupled with blockade of lactate production enhances anti-tumor effects of metformin by boosting T-cell function and limiting cancer cell proliferation [30]. Further, modulation of oxidative phosphorylation (OXPHOS) for overcoming PD-1 resistance to improve anti-tumor responses in specific cancers has been observed [31,32]. These findings suggest that the reprogrammed metabolic axes and enrichment of immune evasion markers can complement each other in driving cancer progression [30,31,33–36]. However, a detailed pan-cancer analysis of such coupling has not yet been conducted.

Here, through a meta-analysis of 184 transcriptomic datasets (Table S1A), and primary tumor samples in multiple carcinomas in The Cancer Genome Atlas (TCGA), and single-cell RNA sequencing data, we evaluate the relationship between these metabolic reprogramming axes and PD-L1 activity and analyze consequences of this association for disease prognosis. We observed a predominant positive correlation of an elevated PD-L1 signature (and CD274 expression levels) with enrichment of partial EMT and glycolysis signatures, but negative correlation with an OXPHOS signatures. Such trends were also largely consistent in a single-cell time-course EMT induction dataset, as well as with other immune checkpoint molecules. Finally, our results revealed that concurrent enrichment of PD-L1 and glycolysis associate with worse patient survival than just glycolysis enrichment without PD-L1, thereby indicating possible synergy in functional aspects of immune suppression and metabolic plasticity in cancer progression.

## Methods

### Software and Datasets

The computational and statistical analysis were conducted using R (version 4.2.1) and Python (version 3.9). Microarray and single-cell RNA sequencing datasets were retrieved from NCBI GEO (Gene expression omnibus) using the ‘GEOquery’ R package. RNA sequencing FASTQ files were obtained from the ENA (European Nucleotide Archive) database. TCGA datasets were obtained using UCSC Xena tools.

### Pre-processing of datasets

The pre-processing of the microarray datasets was conducted to obtain the gene-wise expression from the probe-wise expression matrix using respective annotation files for the mapping of probes to genes. In case multiple probes mapped to a single gene, the mean expression of all mapped probes was utilized to obtain the final values for those genes.

The overall quality of the RNA sequencing datasets was assessed using FastQC. Adapter trimming of FASTQ files was done using ‘Trimmomatic’ (version 0.39) [37] and STAR aligner (version 2.7.10a) [38] was used for the alignment of the reads with hg38-human (or mm10-mouse) reference genome. Raw counts were calculated using HTseq-count and were then normalized for gene length and transformed to TPM (transcripts per million) values. They were then log2 normalized to acquire the final expression data.

For single-cell RNA sequencing datasets, MAGIC (version 2.0.3) [39] imputation algorithm was utilized to recover noisy and sparse single-cell data using diffusion geometry. To map individual reads to particular genes, relevant platform annotation files were utilized.

### EMT scoring methods

For each dataset, several methods were employed to determine EMT scores. Each technique necessitates the input of gene expression data along with a different geneset and algorithm.

### 76GS & KS scores

76GS scores for each sample was calculated using a geneset containing 76 genes [40]. Each sample’s weighted total of the gene expression levels of the 76 genes was calculated, with correlation coefficients to CDH1 expression levels serving as the weighting factors. According to this new scale, a low 76GS score represents a more mesenchymal phenotype whereas a high value indicates predominance of an epithelial one.

The KS technique scores EMT for cell lines and tumor samples using the two-sample Kolmogorov-Smirnov test (KS) [41]. For each of the two signatures (E and M), cumulative distribution functions (CDFs) are produced, and the largest difference between these CDFs is utilized as the test statistic for a two-sample KS test. The resultant EMT scores are in the range [-1, 1]. Mesenchymal and epithelial phenotypes are indicated by positive and negative scores, respectively.

### Epithelial & Mesenchymal scores

To quantify enrichment of epithelial and mesenchymal signatures independently, ssGSEA (single sample gene set enrichment analysis) was performed on KS epithelial (for Epi scores) and KS mesenchymal (for Mes scores) gene signatures separately using GSEAPY python library. The enrichment scores indicate the degree of cordial up/down regulation of genes in the geneset for a given sample. The normalized enrichment score (NES) for these genesets was obtained for further analysis. A higher Epi score denotes a more epithelial phenotype, whereas a higher Mes score signifies enrichment of a mesenchymal phenotype.

### Hallmark EMT & partial EMT scores

ssGSEA method available in the GSEAPY python was used to calculate NES for hallmark EMT geneset from the molecular signatures database (MSigDB) [42]. Partial EMT (pEMT) geneset [43] was utilized to calculate NES for pEMT scores.

### Scoring methods for metabolic pathways and PD-L1

ssGSEA scores were calculated for the hallmark pathway genesets from MSigDB (see supplementary table 1) to obtain the respective normalized gene signature enrichment scores.

AMPK and HIF-1α signatures were quantified using expression levels of their downstream target genes as previously reported [44]. In total, 33 downstream genes for AMPK and 23 downstream genes for HIF-1α were used. FAO2 scores were calculated based on equations previously reported, which use the expression levels of 14 enzyme genes related to FAO [45]. PD-L1 signature was curated as reported earlier [23], wherein the top correlated genes (Spearman correlation coefficient > 0.5 and p < 0.01) with PD-L1 levels in at least any 15 out of the 27 cancer types were considered for this analysis.

The activity scores for metabolic and E/M signatures for the single-cell RNA sequencing datasets were computed using AUCell (version 1.18.1) [46] from the R package ‘Bioconductor’ with default parameters.

### Survival analysis

Survival data (overall survival) were acquired from TCGA. Based on the median of the sample scores, all samples were split into two groups: high PD-L1 and high glycolysis (‘P+G+’) and high PD-L1 but low glycolysis (‘P+G-’). The R package ‘survival’ was employed to perform the Kaplan-Meier analysis, and the plotting was done using ‘ggfortify.’ Reported p-values were calculated using a log rank test. Cox regression was used to determine the hazard ratio (HR) and confidence interval (95% CI) for TCGA cohorts, and forest plots were made using ‘ggplot2’.

## Results

### Enrichment of PD-L1 signature is associated with partial-EMT

To assess the association of PD-L1 with the EMT status of cells, we first calculated single-sample Gene Set Enrichment Analysis (ssGSEA) scores for each sample in 184 bulk transcriptomic datasets (Table S1A) for PD-L1, KS epithelial (Epi), KS mesenchymal (Mes) and partial-EMT gene lists (Table S1B). We also calculated 76GS and KS scores for individual samples in these datasets. Higher Epi and 76GS score signify the enrichment of an epithelial program, whereas a higher KS or Mes score indicates the prevalence of a mesenchymal one.

On correlating the PD-L1 activity signature [23] with Epi scores, a comparable number of datasets showed the correlation to be either positive (n=20; red datapoints) and negative (n=22; blue datapoints) **(Fig 1A)**. On the other hand, PD-L1 activity geneset showed a consistent positive correlation with both Mes and KS scores. Out of 51 datasets that showed significant correlation for PD-L1 vs. Mes scores, 45 of them (88.23%) showed a positive correlation **(Fig 1B)**. Similarly, 73.46% (36 out of 49) datasets showed a positive correlation between KS scores and PD-L1 activity (**Fig 1C)**. Further, PD-L1 scores showed a strong positive skew (41 out of 45 = 91.11%) with the activity of partial EMT geneset [43] **(Fig 1D)**. Together, these results together suggest a link between an enriched PD-L1 signature with a partial EMT program, wherein the cancer cells may acquire a hybrid E/M phenotype, rather than being *completely epithelial or mesenchymal in nature*.

**Figure 1:**
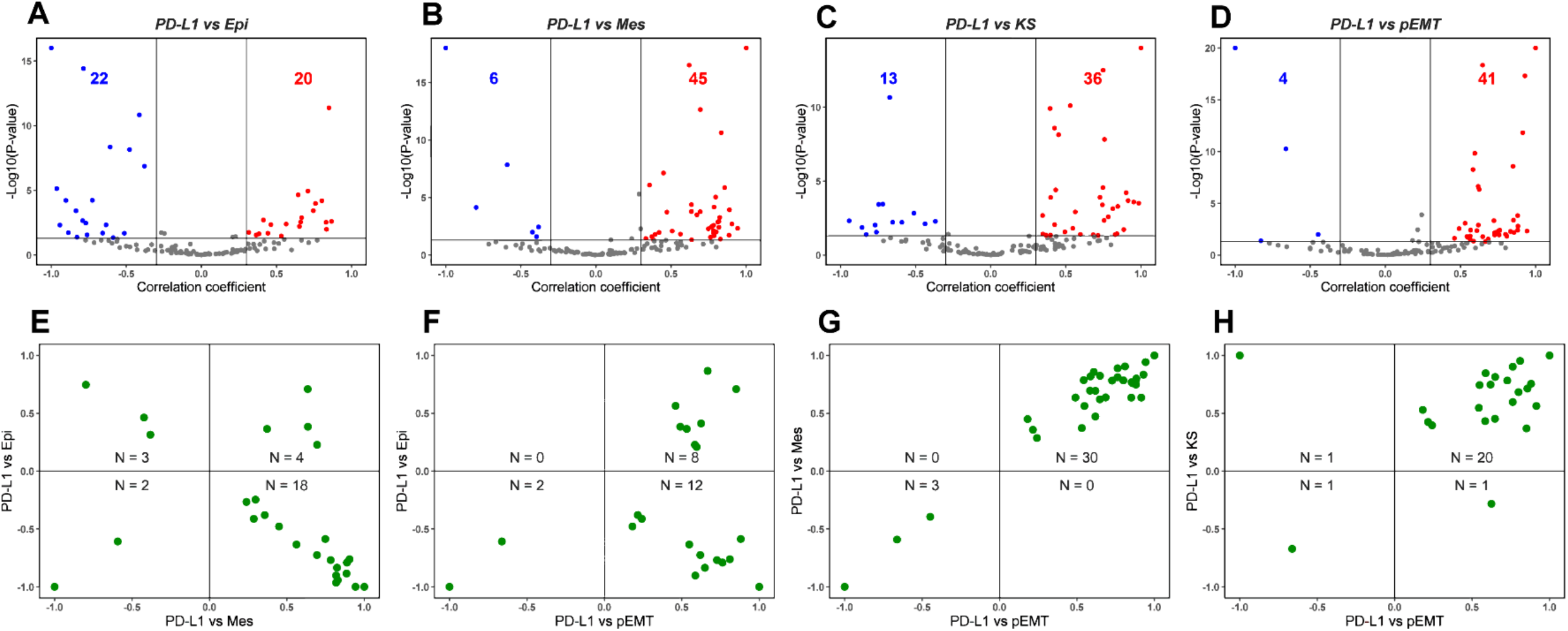
Association between PD-L1 activity gene signature and partial-EMT. **A)** Volcano plots showing correlation coefficient, R (x-axis) and -log_10_(p-value) (y-axis) for PD-L1 vs. Epi scores. Boundaries for significant correlation are set at R > ± 0.3 and p < 0.05. Same as A) but for **B)** PD-L1 vs. Mes, **C)** PD-L1 vs. KS and **D)** PD-L1 vs. pEMT scores. **E)** 2D scatter plot depicting correlation coefficient ‘R’ between PD-L1 vs. Mes (x-axis) and PD-L1 vs. Epi scores (y-axis). ‘N’ denotes the number of significant datapoints lying in each quadrant. Same as E) but for **F)** PD-L1 vs. pEMT and PD-L1 vs. Epi, **G)** PD-L1 vs. pEMT and PD-L1 vs. Mes and **H)** PD-L1 vs. pEMT and PD-L1 vs. KS scores. Each dot denotes a dataset for which this correlation is calculated.

Next, we plotted the correlation coefficients of these pairwise comparisons to further investigate the putative link between PD-L1 and EMT spectrum. In this analysis, we only considered datasets which were significant across both pairwise comparisons. 64.28% (18 out of 28) datasets showed a positive link between PD-L1 and Mes scores while being negatively associated with Epi scores **(Fig 1E)**. In 91.30% (20 out of 22) datasets, PD-L1 activity correlated positively with pEMT scores, among which the correlation with Epi scores can be in either direction to a comparable extent **(Fig 1F)**. Further, among the 33 datasets in which PD-L1 activity associated significantly both with Mes and KS scores, 30 datasets (91.17%) had positive correlation of PD-L1 activity with both the KS and Mes scores **(Fig 1G)**. Reinforcing trends were seen for association of PD-L1 activity with pEMT signature one (**Fig 1H**). Thus, this pan-cancer meta-analysis highlights that the prominent mode of association in these pairwise comparisons is that PD-L1 activity correlates positively with a partial mesenchymal nature, reminiscent of recent experimental reports [19,23,47].

To characterize the connection of immune evasion and EMT extensively, we examined the association of additional immune checkpoint genes with an EMT program. Gene-wise expression values of immune checkpoints - CD274 (encodes PD-L1), CD276, CD47, CTLA4, HAVCR2, LAG3, LGALS9 and PDCD1 were correlated with 76GS, KS and pEMT scores. CD274, CD276 and CD47 correlated predominantly positively with 76GS, KS and pEMT scores **(Fig S1A-C)**, indicating their probable link with partial EMT. Although HAVCR2 and LGALS9 also showed similar trends with pEMT scores, their correlation with 76GS and KS scores did not show a strong skew **(Fig S1E, G)**. Further, CTLA4, PDCD1 and LAG3 had no strong trends in terms of correlation with 76GS, KS or pEMT scores **(Fig S1D, F, H)**. Overall, most immune checkpoint genes associated positively with partial EMT, similar to the trends seen for PD-L1 gene signature.

Together, across this cohort of datasets spanning multiple cancer types (Table S1A), while an enriched PD-L1 gene signature (and gene expression of many immune checkpoints analyzed) did not associate strongly with an epithelial program in either direction (positive or negative), they showed a predominant positive correlation with a mesenchymal signature. This trend was strengthened by the strong positive association of PD-L1 and immune checkpoint genes with the pEMT geneset. Therefore, these results consistently indicate the association of immune evasion with partial EMT program in many carcinomas.

### PD-L1 enrichment is linked to an upregulated glycolysis signature

After examining the association of PD-L1 activity with EMT, we assessed its association with major aspects of energy metabolism that are known to undergo variable degrees of reprogramming during tumor progression, notably Glycolysis and OXPHOS. In the context of cancer progression, several independent studies link upregulation of a glycolytic program and downregulation of OXPHOS with enrichment of PD-L1 [27,33,36]. Consistent with these observations, we observed a strong positive correlation of PD-L1 activity with glycolysis-associated geneset and with HIF-1α, a key glycolytic player. Out of 50 datasets that displayed a significant correlation for the PD-L1 and glycolysis scores, 68% (n=34) datasets show a positive association. PD-L1 activity scores correlate even more strongly with HIF-1α signature, where 38 out of 48 cases (79.17%) reflect a positive correlation between the two. Conversely, among 44 datasets where PD-L1 scores correlated with OXPHOS activity scores, 31 of them (70.45%) exhibited a negative association between the two (**Fig 2A**). These trends were recapitulated in correlation of expression levels of CD274 with the Glycolysis, HIF-1α and OXPHOS signatures (**Fig 2B**). Upon considering pairwise comparisons across individual datasets, glycolysis and HIF-1α enrichment is consistently associated with upregulated PD-L1 signature and CD274 expression **(Fig 2C)**.

**Figure 2:**
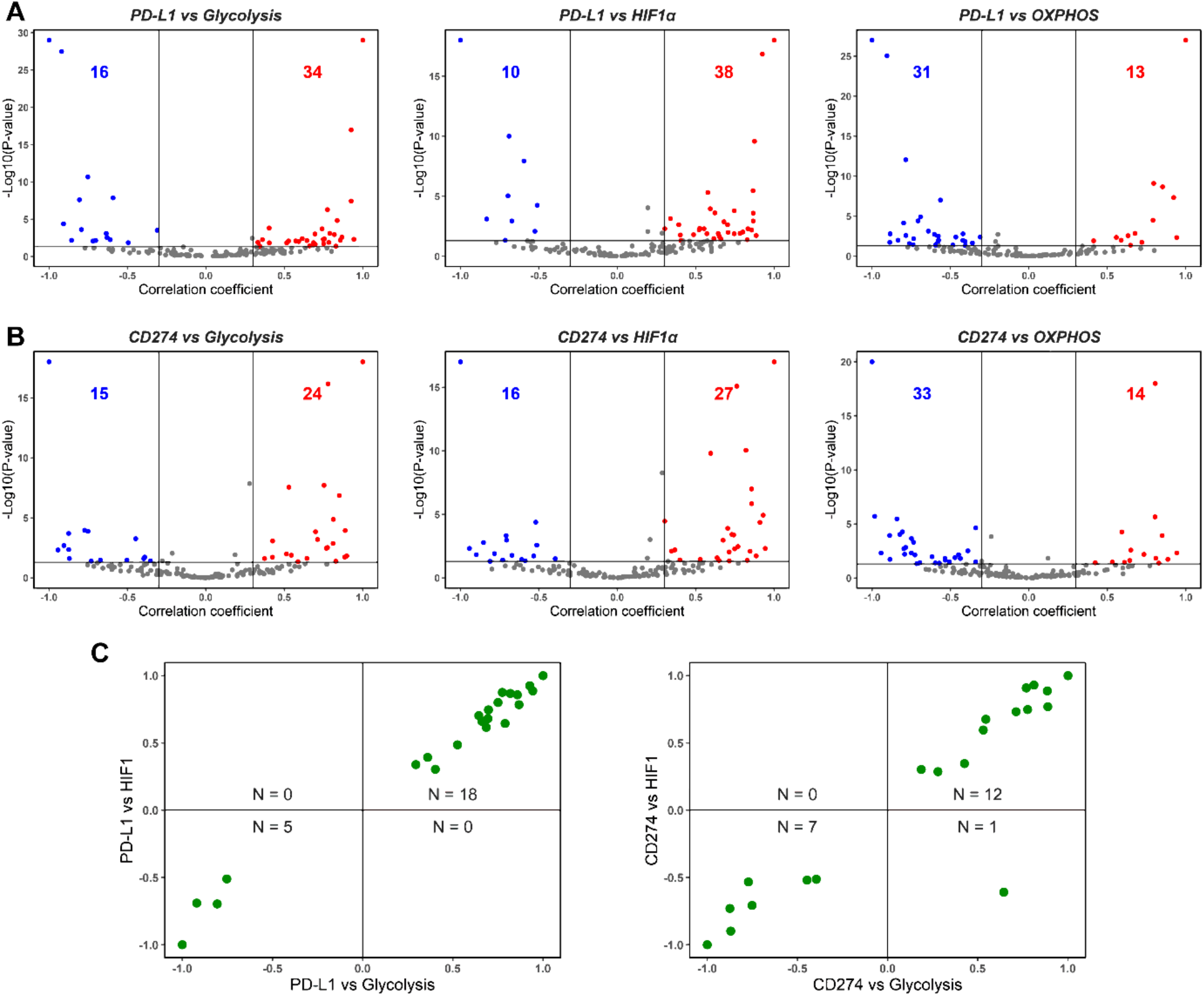
Association between PD-L1, Glycolysis and OXPHOS. **A)** Volcano plots showing correlation coefficient, ‘R’ (x-axis) and -log_10_(p-value) (y-axis) for PD-L1 vs. Glycolysis scores (left) and PD-L1 vs. HIF-1α (middle) and PD-L1 vs. OXPHOS (right). Boundaries for significant correlation are set at R > ± 0.3 and p < 0.05. Same as A) but for **B)** CD274 gene expression vs. Glycolysis (left), CD274 gene expression vs. HIF-1α (middle) and CD274 gene expression vs. OXPHOS (right). **C)** 2D scatter plot depicting correlation coefficient ‘R’ between PD-L1 vs. Glycolysis (x-axis) and PD-L1 vs. HIF-1α scores (y-axis) (left) and CD274 expression vs. Glycolysis (x-axis) and CD274 vs. HIF-1α scores (y-axis) (right). ‘N’ denotes the number of significant datapoints lying in each quadrant.

Among the additional immune checkpoint markers considered here, expression levels of most of them – CD276, CTLA4, HAVCR2, LAG3 and PDCD1 – correlated negatively with OXPHOS. CD47 and CD276 also predominantly exhibited a positive association with HIF1α and/or glycolysis scores (**Fig S2A-B**), while CTLA4, LAG3 and PDCD1 were most likely to be negatively associated with glycolysis (**Fig S2C-G**). These results suggest that glycolysis and OXPHOS may not always be strongly mutually antagonistic to one another, thereby possibly hybrid metabolic (high glycolysis/high OXPHOS) and metabolically quiescent (low glycolysis/low OXPHOS) states, besides the canonical high glycolysis/ low OXPHOS and high OXPHOS/low glycolysis states [48,49].

Overall, both PD-L1 activity scores and CD274 expression levels associate strongly positively with glycolysis and negatively with OXPHOS, consistent with the largely antagonistic trend established for glycolysis and OXPHOS programmes in the context of cancer progression [44]. Further, these strong trends indicate a possible co-operative association between metabolic reprogramming and immune-suppression in a pan-cancer manner.

### Immune checkpoint markers correlate positively with partial EMT, PD-L1 and immune-response signatures in adenocarcinomas

Multiple inflammatory pathways such as tumor necrosis factor-alpha (TNF-α), interferon-gamma (IFN-γ) and nuclear factor-kappa B (NF-κB) are known to induce PD-L1 [50]. Another immune checkpoint marker, LGALS9, was reported to be involved in regulating many of these inflammatory pathways [51]. Thus, we conducted a correlation analysis to elucidate the connection between CD274 (PD-L1 gene) and immune checkpoint markers used earlier (CD47, CD276, CTLA4, HAVCR2, LAG3, LGALS9 and PDCD1). We used well characterized cohorts of primary tumor data from TCGA, in a tissue-specific manner: breast cancer (BRCA), prostate adenocarcinoma (PRAD), bladder cancer (BLCA), stomach adenocarcinomas (STAD), lung adenocarcinoma (LUAD) and pancreatic adenocarcinoma (PAAD).

Across the six abovementioned carcinomas, except occasional deviations seen in PAAD, CD274 expression levels corelated positively with a mesenchymal and a partial EMT signature but correlated negatively with both the hallmark OXPHOS pathway and epithelial signatures (**Fig 3A, left**), consistent with our meta-analysis presented earlier (**Fig 1, 2**). While the association of CD274 expression levels with glycolysis geneset was not as strong across cancer types, they correlated positively with hallmark pathways associated with immune response and inflammation such as INF-α response, IFN-γ response, TNF-α signaling via NF-κB and IL-2/STAT5 signaling. (**Fig 3A, left**).

**Figure 3:**
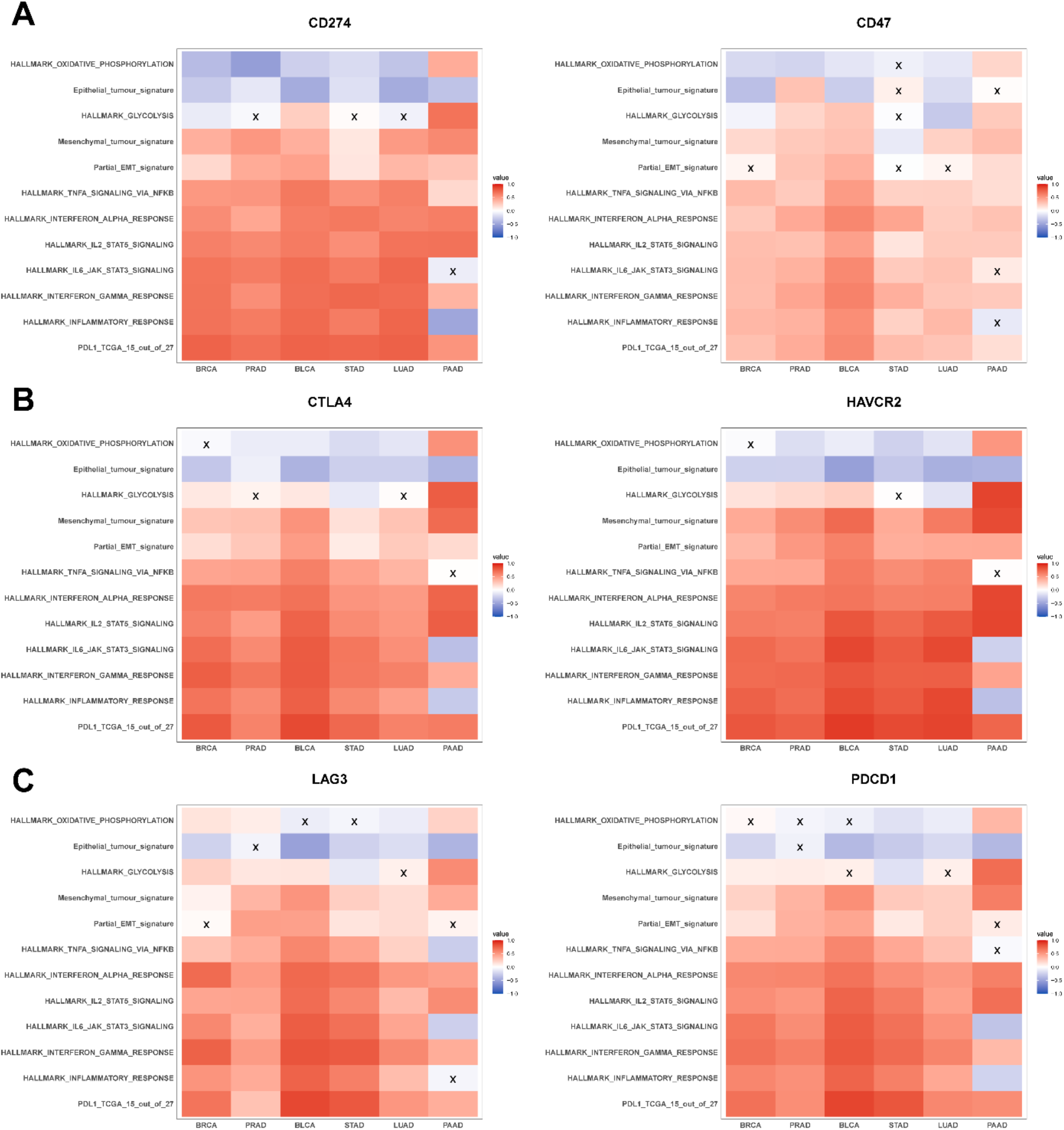
Association of immune checkpoint markers with several hallmark gene signatures in TCGA cohorts: **A)** Heatmap illustrating the spearman correlation coefficients between different hallmark gene signatures with CD274 (left) and CD47 gene expression (right) in BRCA, PRAD, BLCA, STAD, LUAD and PAAD. Insignificant correlations (p >0.05) are marked with ‘X’. Same as A) but for **B)** CTLA4 (left) and HAVCR2 gene expression data (right) and **C)** LAG3 (left) PDCD1 and gene expression data (right).

The abovementioned relationships were largely seen also with the other immune checkpoint genes, where CD47, CTLA4, HAVCR2, LAG3, LGALS9 and PDCD1 correlated positively with metabolic reprogramming and inflammatory signatures, as well as with the mesenchymal and partial EMT ones (**Fig 3A-C, S3A**). However, the negative correlation with epithelial signature was noticed in only 4 out of 8 immune checkpoint genes (**Fig 3, S3A**). Additionally, we noticed that all the immune checkpoint genes (except CD276) were strongly positively correlated with PD-L1 signature, while also positively associating with each other **(Fig 3, S3B)**.

In conclusion, in primary tumor samples in TCGA, multiple immune checkpoint molecules including CD274 correlated positively with inflammatory and immune response associated pathways, PD-L1 signature, and with a mesenchymal behavior. However, the association of these molecules with an epithelial behavior was not as consistently and strongly negative across carcinomas, reminiscent of our previous observations of CD274 expression and PD-L1 signature associating with a partial EMT phenotype. These results augment the trends seen in *in vitro* pan-cancer datasets earlier, establishing a predominant overlap among the enhanced expression of immune checkpoint molecules, partial EMT and metabolic reprogramming aspects.

### Association of CD274 gene expression with partial EMT and metabolic reprogramming is recapitulated in single-cell RNA sequencing data

To further probe the different modalities of association between the EMT status and metabolism reprogramming with PD-L1, we conducted a similar analysis in a single-cell RNA sequencing dataset (GSE147405) [52]. From this dataset, we analyzed the transcriptomic profiles of three cell lines (A549, DU145 and OVCA420) treated with two different EMT-inducers (TGF-β, TNF-α) to analyze how CD274 expression levels change alongside alterations in epithelial-mesenchymal and cellular metabolism status. As a first step, we validated that the EMT scoring metrics used here (KS, Epi, Mes) were consistent in quantifying the E/M status of cells in this dataset **(Fig. S4C)**. While KS score correlated positively with Mes scores consistently across cell lines and treatments, their corelation with Epi scores depended on the cell line – it was expectedly negative in OVCA420, but weakly positive in the two cell lines where Epi and Mes programs were not as strongly antagonistic to one another - A549, DU145. These results further endorse that downregulation of epithelial traits and upregulation of mesenchymal ones may not necessarily happen simultaneously, as often tacitly assumed while claiming EMT [53,54].

In this single-cell dataset, we observed that activity of the mesenchymal geneset correlated positively with CD274 expression across both treatments, further validating the trends observed in bulk datasets **(Fig 4, S4)**. Additionally, CD274 expression leaned strongly towards the positive direction with the epithelial signature in TNF-α-treated DU145 and OVCA420 sample **(Fig 4C, S4B)**, while associating significantly negatively in TGF-β-treated A549 case **(Fig 4A)**. Epi scores were generally non-committed to either direction in the remaining samples **(Fig 4B-C, S4A)**. Apart from TNF-α-treated DU145 sample, in all other cell lines, CD274 correlated positively with pEMT **(Fig 4, S4A-B)**. These results support the observations made in bulk datasets and corroborate the association of CD274 with a partial EMT program.

**Figure 4:**
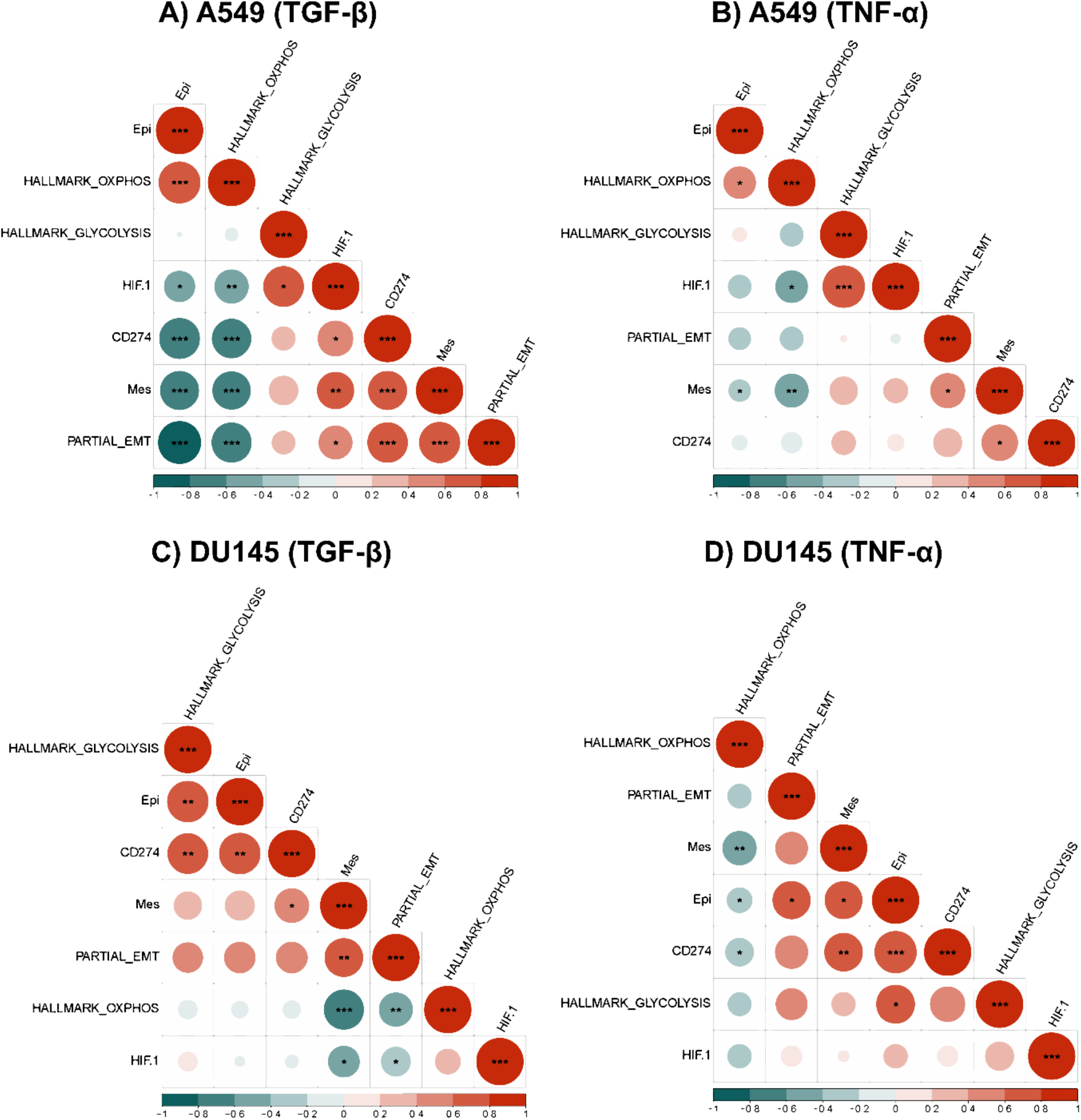
Association between EMT and metabolic axes with CD274 gene expression in single-cell RNA sequencing data (GSE147405). **A)** Heatmap illustrating correlation coefficient ‘R’ for EMT metrics, metabolic pathways and CD274 gene in TGF-β-treated A549 cell line. p-values are calculated using unpaired Students’ T-test with unequal variance and significant correlations are marked with an asterisk (*) for p < 0.05; **: p <0.01; ***: p < 0.001. **B)** Same as A) but for TNF-α-treated A549 cell line. **C)** TGF-β-treated DU145 cell line and **D)** TNF-α-treated DU145 cell line.

We also noticed that in A549 and DU145 cell lines treated with TGF-β and TNF-α, OXPHOS signature consistently displayed a negative correlation with CD274 expression, as seen previously with PD-L1 and OXPHOS in bulk data. Additionally, the glycolysis gene signature revealed a positive correlation between CD274 gene expression levels and PD-L1 gene expression values in bulk data, however this correlation was statistically significant only in TGF-β-treated DU145 cells **(Fig 4C, D)**. On the contrary, in OVCA420 cells treated with TGF-β, both the hallmark OXPHOS and glycolysis signatures showed a significant negative correlation with CD274 expression **(Fig S4A)**, although the negative trend was more pronounced with OXPHOS as compared to glycolysis, showcasing that glycolysis and OXPHOS are not as mutually antagonistic as often presumed. Furthermore, HIF-1α has a significantly positive correlation with CD274 expression.

Therefore, the association of CD274 gene expression with epithelial-hybrid-mesenchymal status of cells is reflected in our analysis of this single-cell dataset, which substantiates previous reports [23] and our analysis of bulk transcriptomic datasets. Similar trends were witnessed for major axes of energy metabolism in cancer cells - glycolysis and OXPHOS, with cell line and/or treatment specific variations altering the extent of correlation seen in this single-cell time-course data [24].

### Survival analysis unfolds the association of concomitant enrichment of PD-L1 and glycolysis with worse patient survival

Finally, we obtained CD274 gene expression and glycolysis scores of patient samples from TCGA across various cancer types to identify any association of metabolic reprogramming and/or immune-suppressive aspects with patient survival. For this assessment, we utilized overall survival (OS) data and for two segregated sample groups, one with a high CD274 and glycolysis score - P+G+ (blue curve in **Fig 5**) and the other with a high CD274 expression but low glycolysis score - P+G- (red curve in **Fig 5**). We observed that P+G+ samples associated with significantly worse patient survival when compared to the P+G-across multiple cancers, indicating that a concurrent upregulation of CD274 expression and glycolysis signatures result in more aggressive disease progression in a pan-cancer manner **(Fig 5)**. Repeating the same analysis with the PD-L1 signature geneset yielded a similar trend, wherein the P+G-samples corresponded with higher OS probability in all TCGA cohorts considered for this analysis **(Fig S6)**.

**Figure 5:**
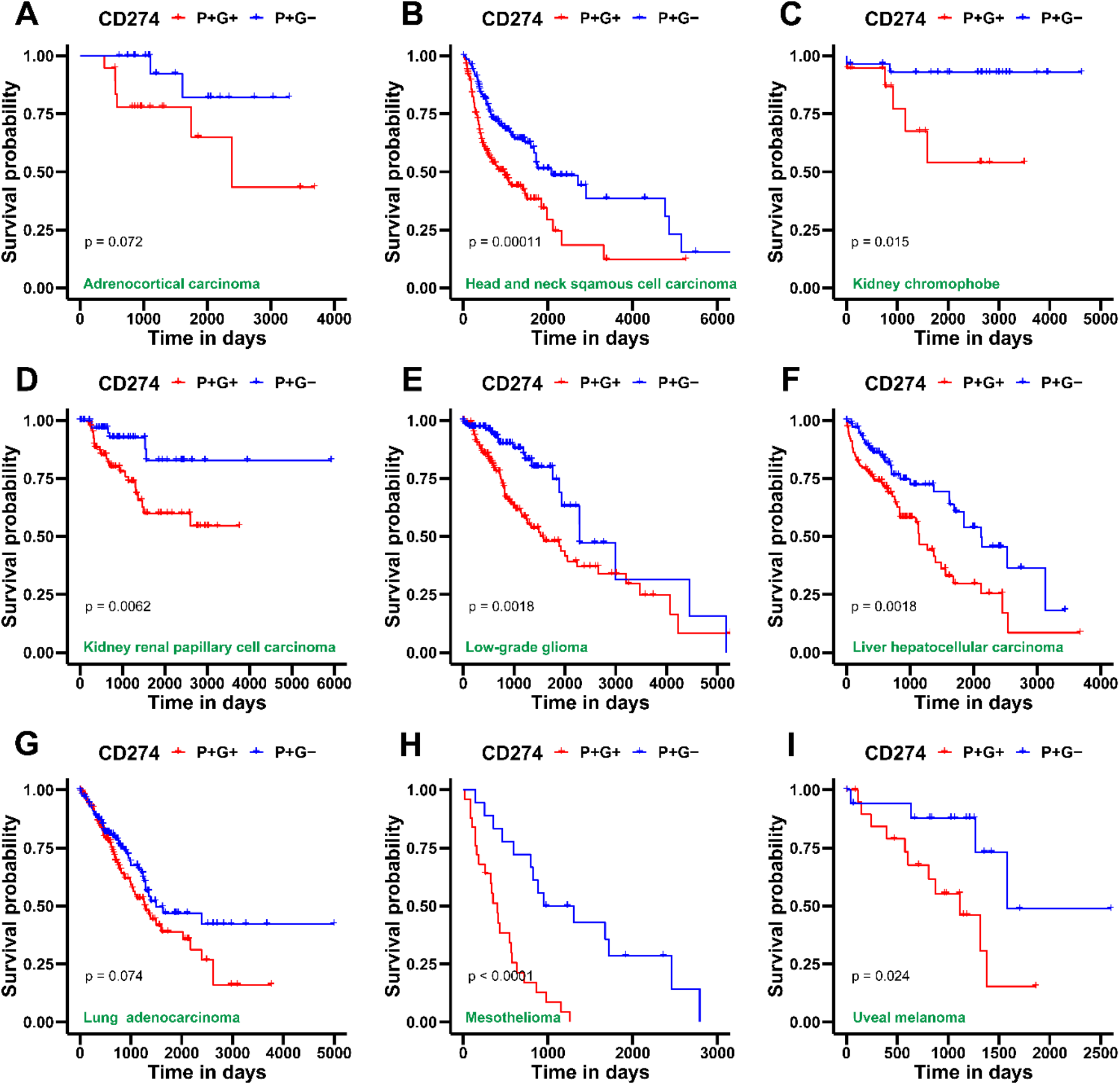
Survival analysis for CD274 gene expression and glycolysis gene signature in several TCGA patient cohorts across cancer types. **A)** Kaplan-Meier curves associating overall survival (OS) with both a high CD274 gene expression and glycolysis signature (blue) and a high CD274 gene expression but a low glycolysis signature (red) in adenoid cystic carcinoma cohort from TCGA. Reported p-values are based on a log-rank test indicating significant difference in survival. Same as A) but for **B)** Head and neck squamous cell carcinoma, **C)** Kidney chromophobe, **D)** Kidney renal papillary cell carcinoma, **E)** Low-grade glioma, **F)** Liver hepatocellular carcinoma, **G)** Lung adenocarcinoma, **H)** Mesothelioma and **I)** Uveal melanoma.

After examining the association of OS data for patient samples with PD-L1 and glycolysis, we investigated the hazard function associated with expression of immune checkpoint markers previously analyzed, in each cancer type in TCGA datasets **(Fig S5)**. Log_2_ hazard ratios (Log_2_HR) corresponding to overall survival for each scenario was calculated. Log_2_HR > 0 indicates an increased risk of morbidity whereas Log_2_HR < 0 signifies better overall survival. This analysis revealed a more context-specific association of the gene expression of immune checkpoint markers with patient outcome. Significant Log_2_HR values reveal the association of higher CD274 expression with worse survival for Skin Cutaneous Melanoma (SKCM), Kidney renal clear cell carcinoma (KIRC) and Adrenocortical carcinoma (ACC), while better patient outcome for Low-Grade Glioma (LGG) **(Fig S5A)**. Elevated CD276 gene expression was predominantly linked with better overall survival **(Fig S5B)**. Log_2_HR ratios were less than zero for CTLA4 gene expression in LGG and KIRC, while being higher than zero in SKCM, Head and Neck Squamous Carcinoma (HNSC) and Breast invasive carcinoma (BRCA) **(Fig S5C)**. Higher HAVCR2 gene expression was associated with lower risk in Uveal Melanoma (UVM), LGG, Thymoma (THYM) and Esophageal carcinoma (ESCA) but higher in Cervical squamous cell carcinoma and endocervical adenocarcinoma (CESC) and SKCM **(Fig. S5D)**. Also, increased LAG3 expression associated with better survival outcome in UVM, LGG and KIRC but worse in SKCM, Thyroid carcinoma (THCA) and Mesothelioma (MESO) **(Fig S5E)**. For high LGALS9 expression, CESC, BRCA, Bladder Urothelial carcinoma (BLCA), HNSC, MESO, Sarcoma (SARC) and SKCM showed worse prognostic probability, while having better patient outcome for UVM, LGG and KIRC **(Fig S5F)**. At last, higher PDCD1 gene expression was linked to lower Log_2_HR in UVM, LGG, Kidney renal papillary cell carcinoma (KIRP) and ESCA, but high Log_2_HR for HNSC, SKCM, Uterine Corpus Endometrial Carcinoma (UCEC) and BRCA **(Fig S5G)**.

In all, these pan-cancer observations offer insights on the classification of patient samples with more vs. less PD-L1 (or CD274) and glycolysis, as well as the worse probability of patient survival linked with their parallel enrichment, which is largely uniform across all evaluated cancer types. In contrast, immune checkpoint gene expression displayed context-dependent relationship with overall survival probabilities for different cancer types.

## Discussion

Tumors often dysregulate the expression of key immune checkpoint proteins [10]. One of such proteins called Programmed Death Ligand-1 (PD-L1) is often employed by cancer cells to bypass the host immunity. The programmed cell death protein-1 (PD-1) on active cytotoxic-T lymphocytes (CTLs) that have infiltrated tumors is detected by PD-L1 on cancer cells and macrophages that effectively turns off their ‘cancer-clearing’ activity through multiple mechanisms. Moreover, when compared to normal tissues, tumor tissues express PD-L1 at considerably higher levels, which drove an interest in inhibiting the PD-L1/PD-1 signaling axis, as an appealing strategy for cancer immunotherapy [55–57]. However, increasing occurrences of resistance in such immunotherapy-based treatments have been reported [58]. It is thus essential to understand the underlying mechanisms behind adaptive immune evasion and its regulation by other molecular factors in the tumor microenvironment (TME) to improve the efficacy of immune checkpoint blockade therapy. In this context, our pan-cancer analysis focusses on specific hallmarks of cancer - EMT and metabolic reprogramming in cancer cells and their association with immune evasion to discern the role of PD-L1 and other checkpoint proteins in cancer progression.

Increasing evidence suggests that PD-L1 regulation is associated with the EMT status of cancer cells. In many carcinomas - BRCA, ESCA and non-small cell lung carcinoma (NSCLC), the EMT status of cells strongly associates with PD-L1 expression levels [59], at least partly through the action of pathways such as phosphoinositide 3-kinase/protein kinase B pathway [60]. Moreover, many EMT-TFs are known modulators of PD-L1 expression, enabling cells in one or more hybrid E/M phenotype(s) to have enriched PD-L1 levels [61,62].

Previous work, including ours, has also indicated association of EMT with metabolic reprogramming [63–65]. Thus, the switching of cancer cell energetics from aerobic respiration (or OXPHOS) to an anaerobic one (or glycolysis) – called as Warburg effect – can impact EMT as well as TME to alter immune-evasive traits. While many cancer cells exhibit a strong propensity towards glycolysis to acclimate themselves with the hypoxic conditions in TME, several studies report that oxidative phosphorylation can remain intact in many different cancers and in a context-specific manner, thus enabling hybrid metabolic phenotypes, akin to the ones reported for EMT extensively now [48]. Such reprogramming can influence both cancer cell behavior and TME to display immunosuppressive characteristics [32,66,67]. A primary reason for this is immunosuppression can be competition in TME brought on by increased glucose demand of cancer cells. Consistently, a study reported that specifically targeting PD-L1 with monoclonal antibodies resulted in decreased glycolysis in tumor cells via obstruction of PI3K/Akt/ mTOR pathway and the translation of glycolytic enzymes, thereby improving the anti-tumor function of T cells [58]. These results reinforce our meta-analysis observations that PD-L1 expression and activity levels are positively correlated with glycolysis pathway in bulk microarray and RNA sequencing datasets, with cancer-specific differences shaping the trends in TCGA cohort and single-cell analysis. Our observations on correlation between HIF-1α and PD-L1 resonate with experimental observations in tumor-bearing mice reporting that PD-L1 upregulation in hypoxia depended on HIF-1α activity [68].

Although our analysis suggests possible synergy among the different hallmarks of tumor progression, we did not perform any specific analysis to elucidate the direction of mechanistic influence of these cellular programs precisely. Thus, a causal relationship among these axes needs to be yet elucidated through specific perturbation experiments. Such patterns of association could help in developing more effective anti-tumor effective strategies in future to tackle the clinical challenges of tumor plasticity and heterogeneity that tend to improve the fitness of cancer often as a whole [69].

## Author contributions

MKJ designed and supervised research. SM, MS and SR performed research and contributed to manuscript writing. SM (Susmita Mandal), SSM, YM, AS performed research. All authors contributed to revising the manuscript draft.

## Supplementary Figures

**Figure S1:**
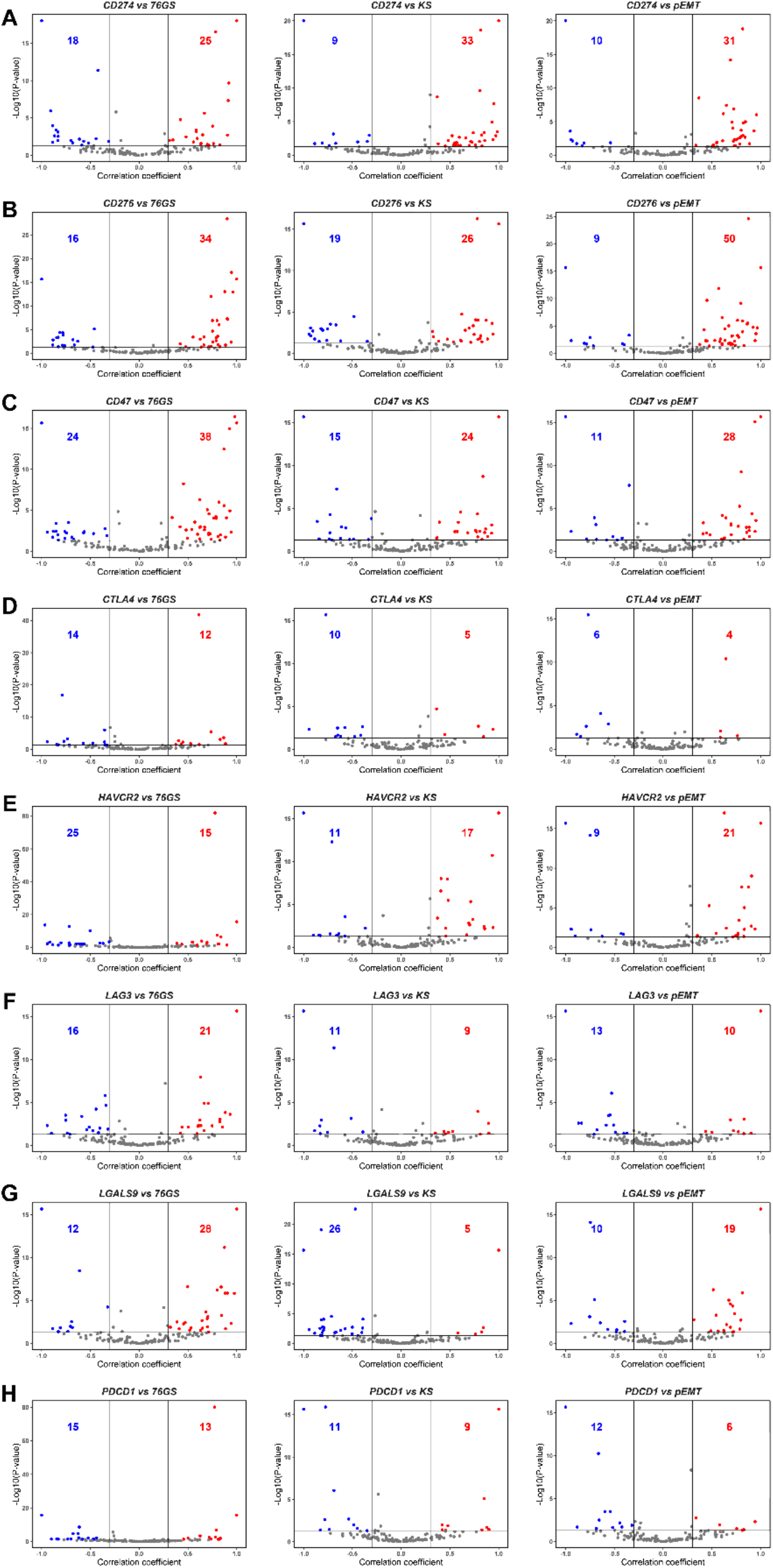
Relationship of immune checkpoint markers with EMT in bulk transcriptomics datasets. **A)** Volcano plots showing correlation coefficient, ‘R’ (x-axis) and -log_10_(p-value) (y-axis) for CD274 gene expression vs. 76GS (left), CD274 vs. KS (middle) and CD274 vs. pEMT scores (right). Boundaries for significant correlation are set at R > ± 0.3 and p < 0.05. Same as A) but for **B)** CD276 gene expression vs. 76GS (left), CD276 vs. KS (middle) and CD276 vs. pEMT scores (right), **C)** CD47 gene expression vs. 76GS (left), CD47 vs. KS (middle) and CD47 vs. pEMT scores (right), **D)** CTLA4 gene expression vs. 76GS (left), CTLA4 vs. KS (middle) and CTLA4 vs. pEMT scores (right), **E)** HAVCR2 gene expression vs. 76GS (left), HAVCR2 vs. KS (middle) and HAVCR2 vs. pEMT scores (right), **F)** LAG3 gene expression vs. 76GS (left), LAG3 vs. KS (middle) and LAG3 vs. pEMT scores (right), **G)** LGALS9 gene expression vs. 76GS (left), LGALS9 vs. KS (middle) and LGALS9 vs. pEMT scores (right), and **H)** PDCD1 gene expression vs. 76GS (left), PDCD1 vs. KS (middle) and PDCD1 vs. pEMT scores (right).

**Figure S2:**
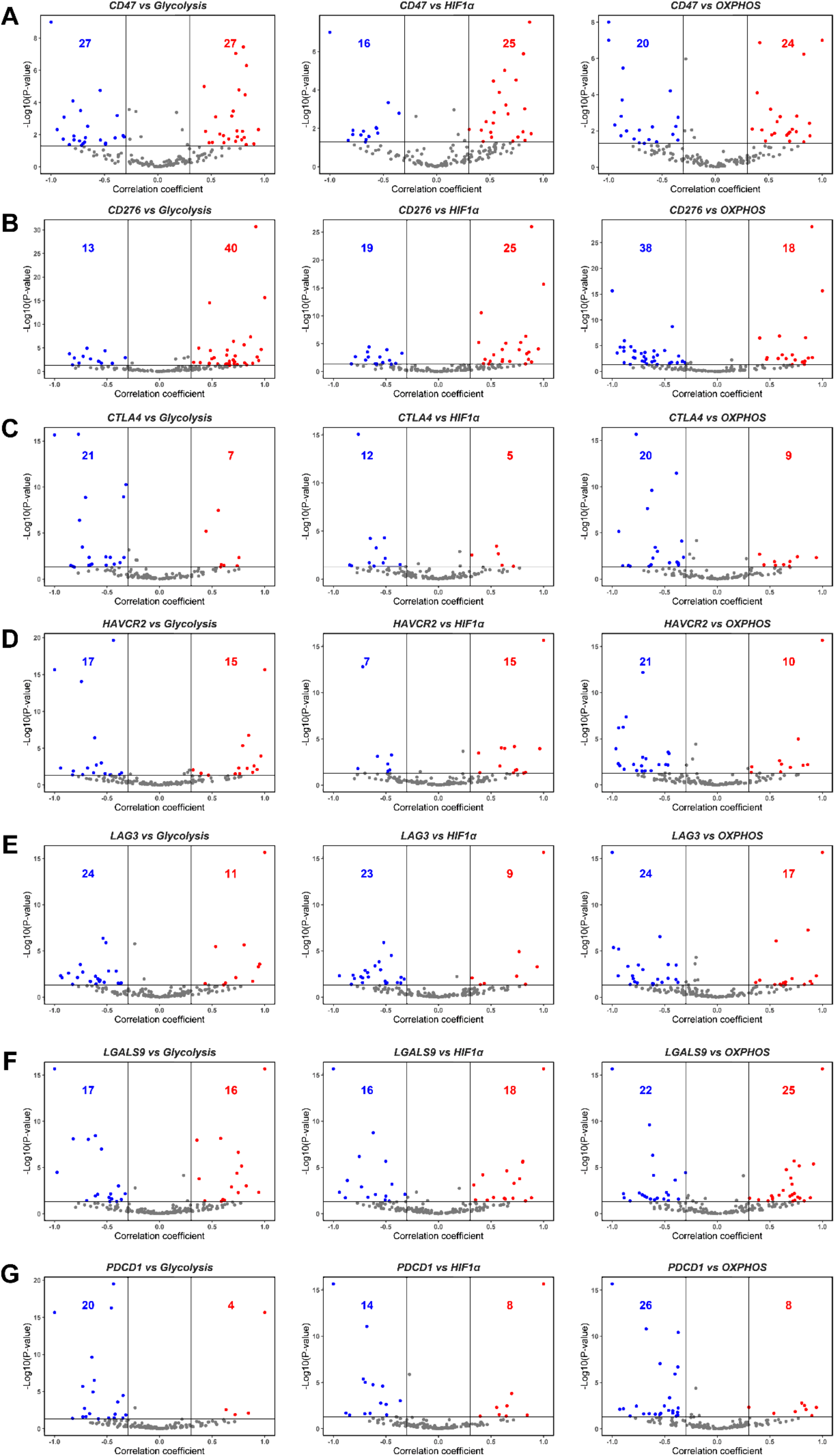
Relationship of immune checkpoint markers with glycolysis (its master regulator HIF-1α) and OXPHOS gene signatures. **A)** Volcano plots showing correlation coefficient, ‘R’ (x-axis) and - log_10_(p-value) (y-axis) for CD47 gene expression vs. glycolysis (left), CD47 vs. HIF1-α (middle) and CD47 vs. OXPHOS scores (right). Boundaries for significant correlation are set at R > ± 0.3 and p < 0.05. Same as A) but for **B)** CD276 gene expression vs. glycolysis (left), CD276 vs. HIF1-α (middle) and CD276 vs. OXPHOS scores (right), **C)** CTLA4 gene expression vs. glycolysis (left), CTLA4 vs. HIF1-α (middle) and CTLA4 vs. OXPHOS scores (right), **D)** HAVCR2 gene expression vs. glycolysis (left), HAVCR2 vs. HIF1-α (middle) and HAVCR2 vs. OXPHOS scores (right), **E)** LAG3 gene expression vs. glycolysis (left), LAG3 vs. HIF1-α (middle) and LAG3 vs. OXPHOS scores (right), **F)** LGALS9 gene expression vs. glycolysis (left), LGALS9 vs. HIF1-α (middle) and LGALS9 vs. OXPHOS scores (right) and **G)** PDCD1 gene expression vs. glycolysis (left), PDCD1 vs. HIF1-α (middle) and PDCD1 vs. OXPHOS scores (right).

**Figure S3:**
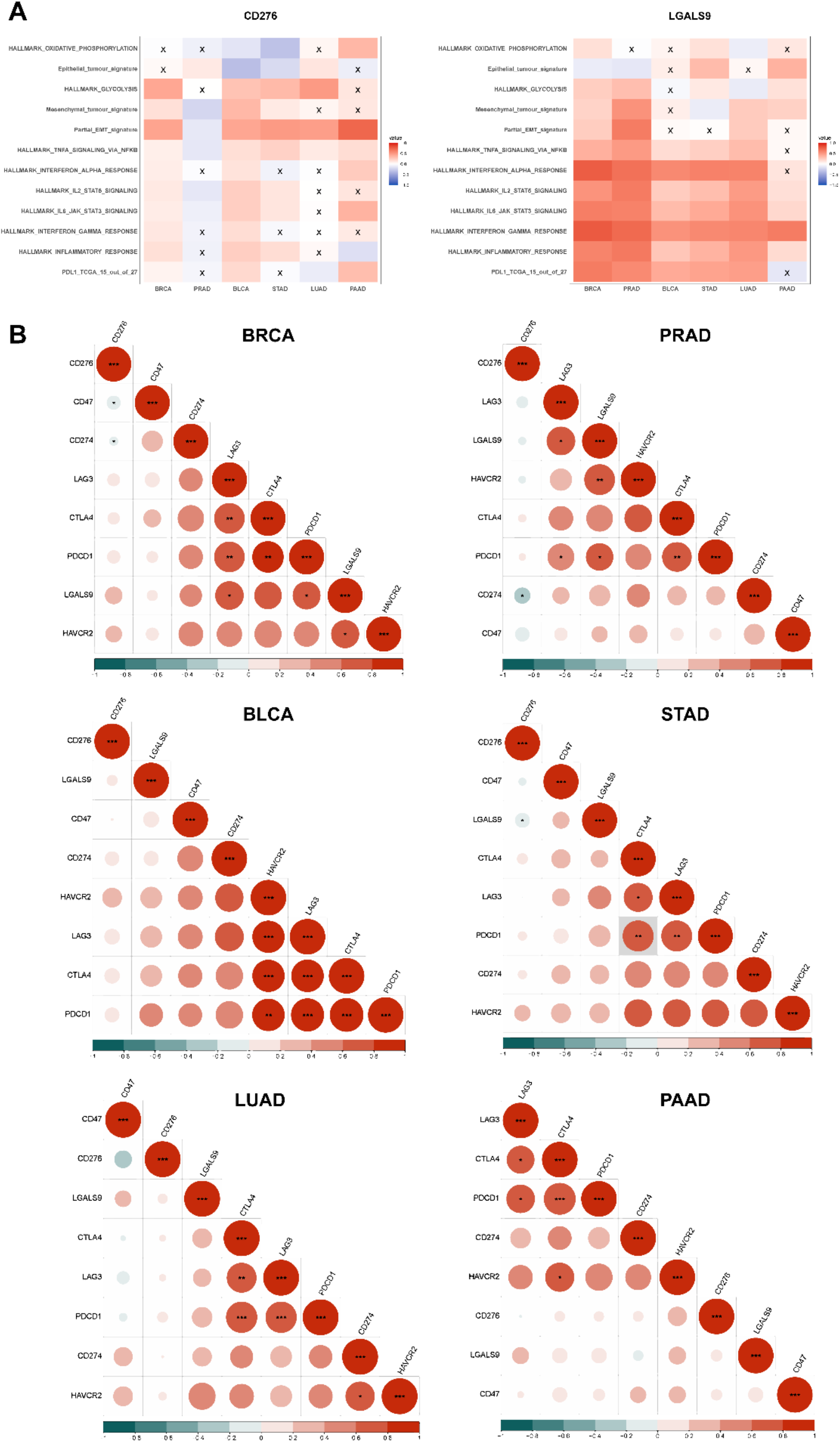
Association of immune checkpoint markers with hallmark pathways in TCGA datasets. **A)** Heatmap illustrating the spearman correlation coefficients between different hallmark gene signatures with CD276 (left) and LGALS9 gene expression (right) in BRCA, PRAD, BLCA, STAD, LUAD and PAAD. Insignificant correlations (p >0.05) are marked with ‘X’. **B)** Heatmap showing correlation between different immune checkpoints in BRCA, PRAD, BLCA, STAD, LUAD and PAAD. p-values are calculated using unpaired Students’ T-test with unequal variance and significant correlations are marked with an asterisk (*) for p < 0.05; **: p <0.01; ***: p < 0.001.

**Figure S4:**
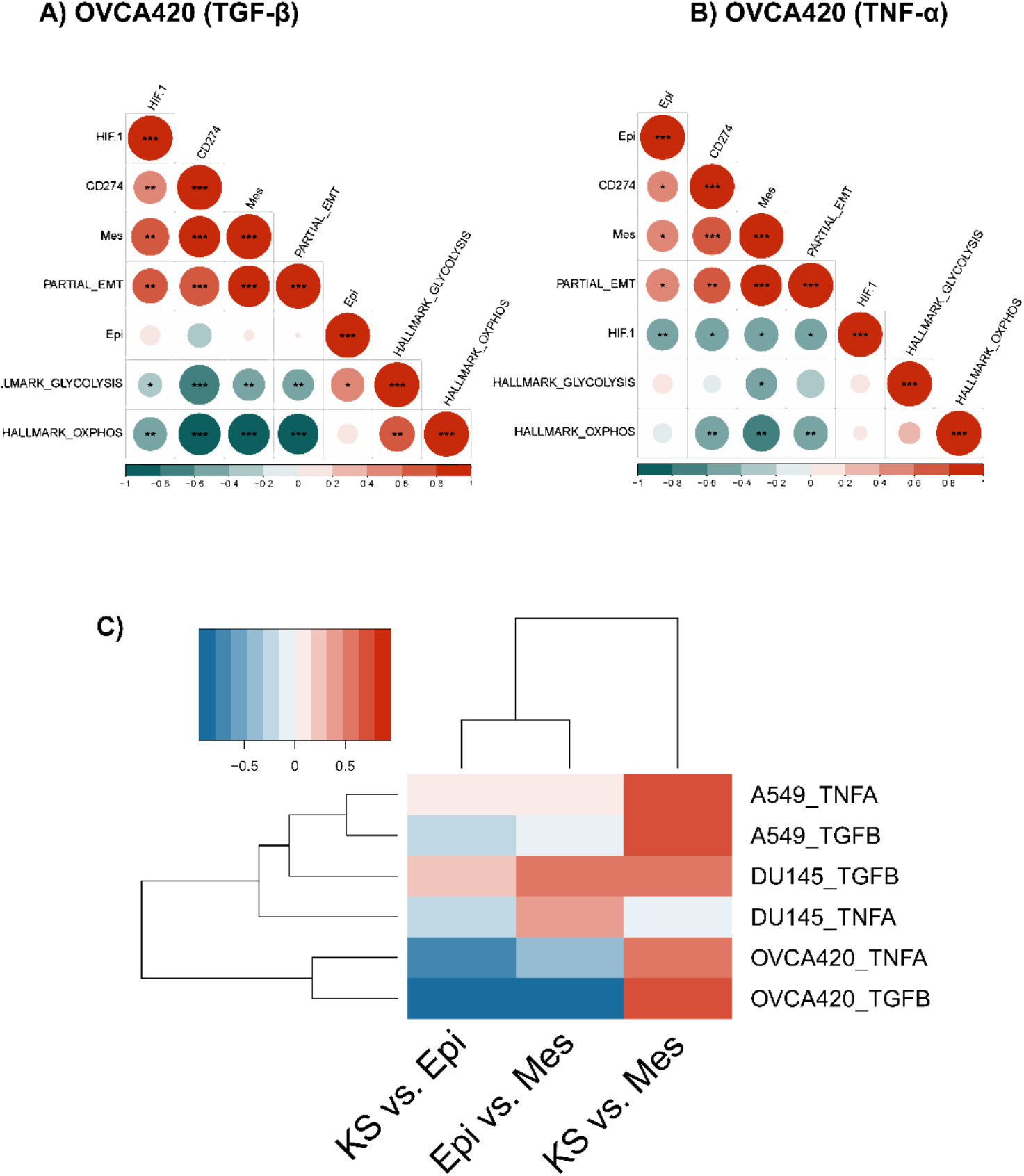
Association between EMT and metabolic axes with CD274 gene expression in single-cell RNA sequencing data (GSE147405). **A)** Heatmap illustrating correlation coefficient ‘R’ for EMT metrics, metabolic pathways and CD274 gene in TGF-β-treated OVCA420 cell line. p-values are calculated using unpaired Students’ T-test with unequal variance and significant correlations are marked with an asterisk (*) for p < 0.05; **: p <0.01; ***: p < 0.001. **B)** Same as A) but for TNF-α-treated OVCA420 cell line. **C)** Heatmap displaying spearman’s correlation coefficient for KS vs. Epi, Epi vs. Mes, and KS vs. Mes scores in A549, DU145 and OVCA420 cell lines across both treatments. All depicted correlations are significant.

**Figure S5:**
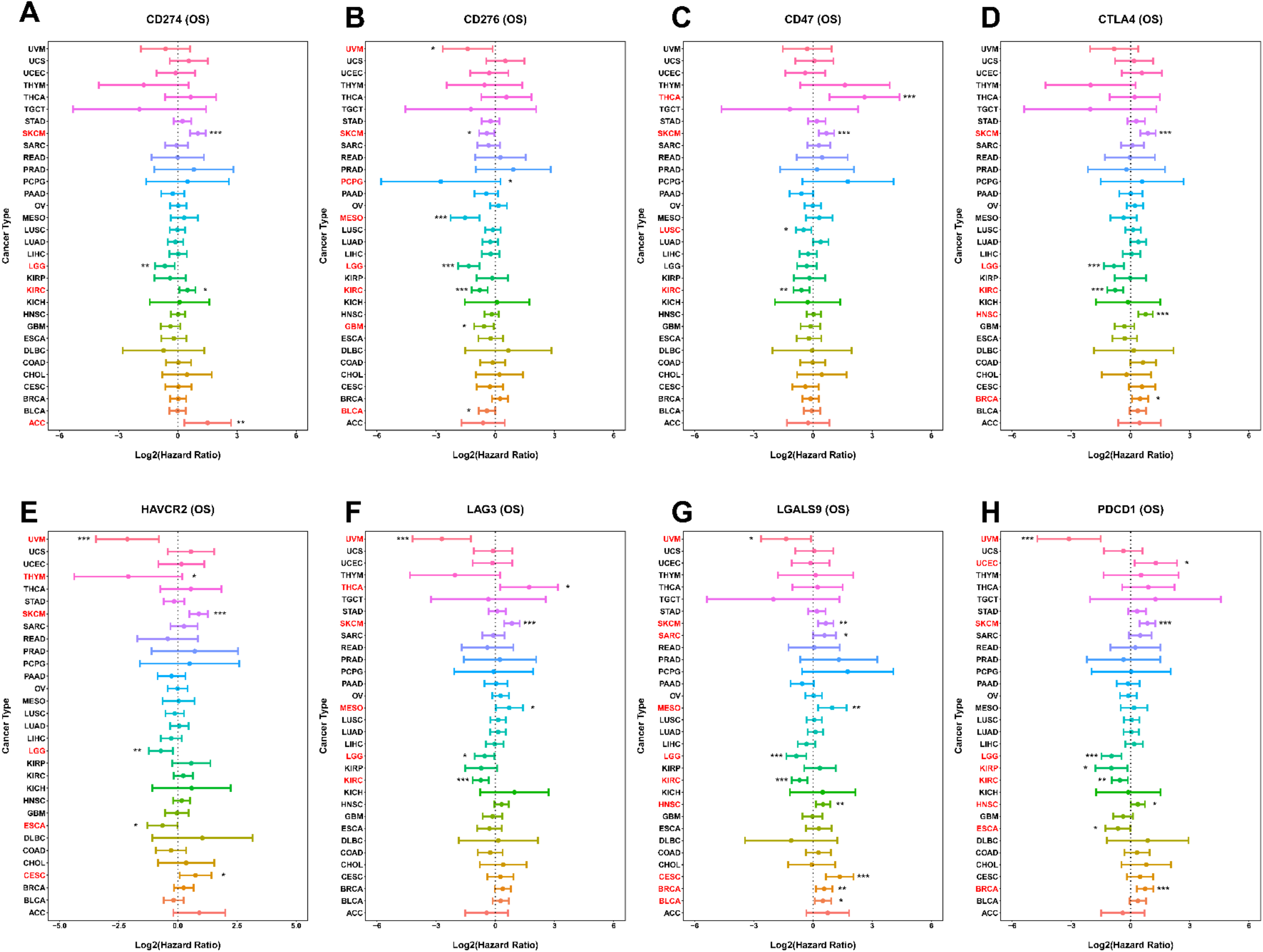
Cox proportional hazard ratios for immune checkpoint markers in different TCGA datasets. **A)** Forest plot depicting Log_2_ hazard ratios (HR; mean ± 95% confidence interval) for overall survival associated with CD274 gene expression across several TCGA cancer types. p-values are based on log-rank test and significant difference in survival are marked with an asterisk (*) for p < 0.05; **: p <0.01; ***: p < 0.001. Axis titles of cancer types with significant differences (p < 0.05) are also highlighted in red, while insignificant ones are shown in black. Same as A) but for **B)** CD276, **C)** CD47, **D)** CTLA4, **E)** HAVCR2, **F)** LAG3, **G)** LGALS9 and **H)** PDCD1 gene expression.

**Figure S6:**
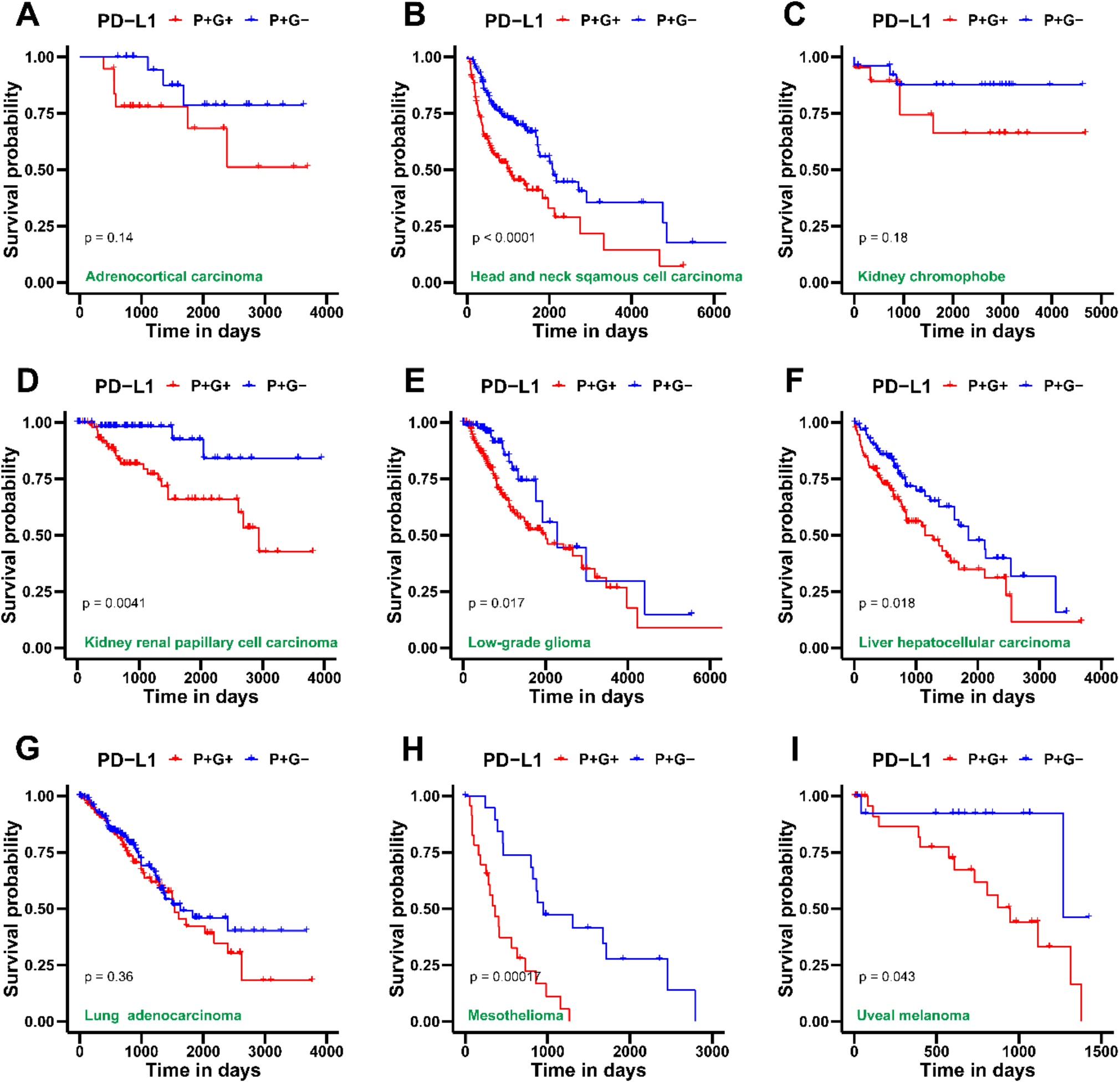
Survival analysis for PD-L1 and glycolysis gene signatures in several TCGA patient cohorts across cancer types. **A)** Kaplan-Meier curves associating overall survival (OS) with both a high PD-L1 and glycolysis signature (blue) and an enriched PD-L1 but a low glycolysis signature (red) in adenoid cystic carcinoma cohort from TCGA. Reported p-values are based on a log-rank test indicating significant difference in survival. Same as A) but for **B)** Head and neck squamous cell carcinoma, **C)** Kidney chromophobe, **D)** Kidney renal papillary cell carcinoma, **E)** Low-grade glioma, **F)** Liver hepatocellular carcinoma, **G)** Lung adenocarcinoma, **H)** Mesothelioma and **I)** Uveal melanoma.

## Supplementary Table Legends

***Supplementary Table 1A:*** *Description of the 184 bulk transcriptomic datasets used in this study*.

***Supplementary Table 1B:*** *Genesets used for scoring EMT, metabolic and PD-L1 gene signatures*.

***Supplementary Table 1C:*** *Spearman’s Correlation coefficient ‘R’ and corresponding p-values for correlation of PD-L1 gene signature and immune checkpoint markers with KS, 76GS, Epi, Mes and pEMT scores*.

***Supplementary Table 2:*** *Spearman’s Correlation coefficient ‘R’ and corresponding p-values for correlation of PD-L1 gene signature and immune checkpoint markers with glycolysis, HIF-1α and OXPHOS scores*.

***Supplementary Table 3:*** *Spearman’s Correlation coefficient ‘R’ and corresponding p-values for correlation of immune checkpoint markers with different hallmark pathways in TCGA cohorts*.

***Supplementary Table 4A:*** *Spearman’s Correlation coefficient ‘R’ and corresponding p-values for correlation between CD274 gene expression, E/M scores, and scores for metabolism in cell line samples of GSE147405*.

***Supplementary Table 4B:*** *Spearman’s Correlation coefficient ‘R’ and corresponding p-values for correlation between KS vs. Mes, Epi vs. Mes and Ks vs. Epi for cell line samples in GSE147405*.

***Supplementary Table 5:*** *Log2 hazard ratios, mean ± 95% confidence intervals and corresponding p-values for overall survival associated with CD274 gene expression across several TCGA cancer types*.

## References

[1] D. Hanahan, R.A. Weinberg, Hallmarks of Cancer: The Next Generation, Cell. 144 (2011) 646–674. https://doi.org/10.1016/j.cell.2011.02.013.

[2] D.S. Vinay, E.P. Ryan, G. Pawelec, W.H. Talib, J. Stagg, E. Elkord, T. Lichtor, W.K. Decker, R.L. Whelan, H.M.C.S. Kumara, E. Signori, K. Honoki, A.G. Georgakilas, A. Amin, W.G. Helferich, C.S. Boosani, G. Guha, M.R. Ciriolo, S. Chen, S.I. Mohammed, A.S. Azmi, W.N. Keith, A. Bilsland, D. Bhakta, D. Halicka, H. Fujii, K. Aquilano, S.S. Ashraf, S. Nowsheen, X. Yang, B.K. Choi, B.S. Kwon, Immune evasion in cancer: Mechanistic basis and therapeutic strategies, Semin Cancer Biol. 35 (2015) S185–S198. https://doi.org/10.1016/j.semcancer.2015.03.004.

[3] D.M. Pardoll, The blockade of immune checkpoints in cancer immunotherapy, Nat Rev Cancer. 12 (2012) 252–264. https://doi.org/10.1038/nrc3239.

[4] J.C. Becker, M.H. Andersen, D. Schrama, P. thor Straten, Immune-suppressive properties of the tumor microenvironment, Cancer Immunology, Immunotherapy. 62 (2013) 1137– 1148. https://doi.org/10.1007/s00262-013-1434-6.

[5] G.J. Freeman, A.J. Long, Y. Iwai, K. Bourque, T. Chernova, H. Nishimura, L.J. Fitz, N. Malenkovich, T. Okazaki, M.C. Byrne, H.F. Horton, L. Fouser, L. Carter, V. Ling, M.R. Bowman, B.M. Carreno, M. Collins, C.R. Wood, T. Honjo, Engagement of the Pd-1 Immunoinhibitory Receptor by a Novel B7 Family Member Leads to Negative Regulation of Lymphocyte Activation, Journal of Experimental Medicine. 192 (2000) 1027–1034. https://doi.org/10.1084/jem.192.7.1027.

[6] P. Kolar, K. Knieke, J.K.E. Hegel, D. Quandt, G.-R. Burmester, H. Hoff, M.C. Brunner-Weinzierl, CTLA-4 (CD152) controls homeostasis and suppressive capacity of regulatory T cells in mice, Arthritis Rheum. 60 (2009) 123–132. https://doi.org/10.1002/art.24181.

[7] J. Leitner, C. Klauser, W.F. Pickl, J. Stöckl, O. Majdic, A.F. Bardet, D.P. Kreil, C. Dong, T. Yamazaki, G. Zlabinger, K. Pfistershammer, P. Steinberger, B7-H3 is a potent inhibitor of human T-cell activation: No evidence for B7-H3 and TREML2 interaction, Eur J Immunol. 39 (2009) 1754–1764. https://doi.org/10.1002/eji.200839028.

[8] J.F. Grosso, C.C. Kelleher, T.J. Harris, C.H. Maris, E.L. Hipkiss, A. de Marzo, R. Anders, G. Netto, D. Getnet, T.C. Bruno, M. v. Goldberg, D.M. Pardoll, C.G. Drake, LAG-3 regulates CD8+ T cell accumulation and effector function in murine self- and tumor-tolerance systems, Journal of Clinical Investigation. 117 (2007) 3383–3392. https://doi.org/10.1172/JCI31184.

[9] H. Li, D. Yang, M. Hao, H. Liu, Differential expression of HAVCR2 gene in pan-cancer: A potential biomarker for survival and immunotherapy, Front Genet. 13 (2022). https://doi.org/10.3389/fgene.2022.972664.

[10] D.M. Pardoll, The blockade of immune checkpoints in cancer immunotherapy, Nat Rev Cancer. 12 (2012) 252–264. https://doi.org/10.1038/nrc3239.

[11] C. Sun, R. Mezzadra, T.N. Schumacher, Regulation and Function of the PD-L1 Checkpoint, Immunity. 48 (2018) 434–452. https://doi.org/10.1016/j.immuni.2018.03.014.

[12] M.J. Butte, M.E. Keir, T.B. Phamduy, A.H. Sharpe, G.J. Freeman, Programmed Death-1 Ligand 1 Interacts Specifically with the B7-1 Costimulatory Molecule to Inhibit T Cell Responses, Immunity. 27 (2007) 111–122. https://doi.org/10.1016/j.immuni.2007.05.016.

[13] S. Munir, G.H. Andersen, Ö. Met, M. Donia, T.M. Frøsig, S.K. Larsen, T.W. Klausen, I.M. Svane, M.H. Andersen, HLA-Restricted CTL That Are Specific for the Immune Checkpoint Ligand PD-L1 Occur with High Frequency in Cancer Patients, Cancer Res. 73 (2013) 1764–1776. https://doi.org/10.1158/0008-5472.CAN-12-3507.

[14] M.W.L. Teng, S.F. Ngiow, A. Ribas, M.J. Smyth, Classifying Cancers Based on T-cell Infiltration and PD-L1, Cancer Res. 75 (2015) 2139–2145. https://doi.org/10.1158/0008-5472.CAN-15-0255.

[15] T. Maeda, M. Hiraki, C. Jin, H. Rajabi, A. Tagde, M. Alam, A. Bouillez, X. Hu, Y. Suzuki, M. Miyo, T. Hata, K. Hinohara, D. Kufe, MUC1-C Induces PD-L1 and Immune Evasion in Triple-Negative Breast Cancer, Cancer Res. 78 (2018) 205–215. https://doi.org/10.1158/0008-5472.CAN-17-1636.

[16] G. Ma, Y. Liang, Y. Chen, L. Wang, D. Li, Z. Liang, X. Wang, D. Tian, X. Yang, H. Niu, Glutamine Deprivation Induces PD-L1 Expression via Activation of EGFR/ERK/c-Jun Signaling in Renal Cancer, Molecular Cancer Research. 18 (2020) 324–339. https://doi.org/10.1158/1541-7786.MCR-19-0517.

[17] C. Sun, R. Mezzadra, T.N. Schumacher, Regulation and Function of the PD-L1 Checkpoint, Immunity. 48 (2018) 434–452. https://doi.org/10.1016/j.immuni.2018.03.014.

[18] C.-Y. Ock, S. Kim, B. Keam, M. Kim, T.M. Kim, J.-H. Kim, Y.K. Jeon, J.-S. Lee, S.K. Kwon, J.H. Hah, T.-K. Kwon, D.-W. Kim, H.-G. Wu, M.-W. Sung, D.S. Heo, PD-L1 expression is associated with epithelial-mesenchymal transition in head and neck squamous cell carcinoma, Oncotarget. 7 (2016) 15901–15914. https://doi.org/10.18632/oncotarget.7431.

[19] A. Dongre, M. Rashidian, F. Reinhardt, A. Bagnato, Z. Keckesova, H.L. Ploegh, R.A. Weinberg, Epithelial-to-Mesenchymal Transition Contributes to Immunosuppression in Breast Carcinomas, Cancer Res. 77 (2017) 3982–3989. https://doi.org/10.1158/0008-5472.CAN-16-3292.

[20] A. Asgarova, K. Asgarov, Y. Godet, P. Peixoto, A. Nadaradjane, M. Boyer-Guittaut, J. Galaine, D. Guenat, V. Mougey, J. Perrard, J.R. Pallandre, A. Bouard, J. Balland, C. Tirole, O. Adotevi, E. Hendrick, M. Herfs, P.F. Cartron, C. Borg, E. Hervouet, PD-L1 expression is regulated by both DNA methylation and NF-kB during EMT signaling in non-small cell lung carcinoma, Oncoimmunology. 7 (2018) e1423170. https://doi.org/10.1080/2162402X.2017.1423170.

[21] D. Imai, T. Yoshizumi, S. Okano, S. Itoh, T. Ikegami, N. Harada, S. Aishima, Y. Oda, Y. Maehara, IFN-γ Promotes Epithelial-Mesenchymal Transition and the Expression of PD-L1 in Pancreatic Cancer, Journal of Surgical Research. 240 (2019) 115–123. https://doi.org/10.1016/j.jss.2019.02.038.

[22] L. Chen, D.L. Gibbons, S. Goswami, M.A. Cortez, Y.-H. Ahn, L.A. Byers, X. Zhang, X. Yi, D. Dwyer, W. Lin, L. Diao, J. Wang, J.D. Roybal, M. Patel, C. Ungewiss, D. Peng, S. Antonia, M. Mediavilla-Varela, G. Robertson, S. Jones, M. Suraokar, J.W. Welsh, B. Erez, I.I. Wistuba, L. Chen, D. Peng, S. Wang, S.E. Ullrich, J. v. Heymach, J.M. Kurie, F.X.-F. Qin, Metastasis is regulated via microRNA-200/ZEB1 axis control of tumour cell PD-L1 expression and intratumoral immunosuppression, Nat Commun. 5 (2014) 5241. https://doi.org/10.1038/ncomms6241.

[23] S. Sahoo, S.P. Nayak, K. Hari, P. Purkait, S. Mandal, A. Kishore, H. Levine, M.K. Jolly, Immunosuppressive Traits of the Hybrid Epithelial/Mesenchymal Phenotype, Front Immunol. 12 (2021). https://doi.org/10.3389/fimmu.2021.797261.

[24] S. Muralidharan, S. Sahoo, A. Saha, S. Chandran, S.S. Majumdar, S. Mandal, H. Levine, M.K. Jolly, Quantifying the Patterns of Metabolic Plasticity and Heterogeneity along the Epithelial–Hybrid–Mesenchymal Spectrum in Cancer, Biomolecules. 12 (2022) 297. https://doi.org/10.3390/biom12020297.

[25] I. Georgakopoulos-Soares, D. v. Chartoumpekis, V. Kyriazopoulou, A. Zaravinos, EMT Factors and Metabolic Pathways in Cancer, Front Oncol. 10 (2020). https://doi.org/10.3389/fonc.2020.00499.

[26] M. Sciacovelli, C. Frezza, Metabolic reprogramming and epithelial-to-mesenchymal transition in cancer, FEBS J. 284 (2017) 3132–3144. https://doi.org/10.1111/febs.14090.

[27] I. Kareva, P. Hahnfeldt, The Emerging “Hallmarks” of Metabolic Reprogramming and Immune Evasion: Distinct or Linked?, Cancer Res. 73 (2013) 2737–2742. https://doi.org/10.1158/0008-5472.CAN-12-3696.

[28] C.-H. Chang, J. Qiu, D. O’Sullivan, M.D. Buck, T. Noguchi, J.D. Curtis, Q. Chen, M. Gindin, M.M. Gubin, G.J.W. van der Windt, E. Tonc, R.D. Schreiber, E.J. Pearce, E.L. Pearce, Metabolic Competition in the Tumor Microenvironment Is a Driver of Cancer Progression, Cell. 162 (2015) 1229–1241. https://doi.org/10.1016/j.cell.2015.08.016.

[29] K. Fischer, P. Hoffmann, S. Voelkl, N. Meidenbauer, J. Ammer, M. Edinger, E. Gottfried, S. Schwarz, G. Rothe, S. Hoves, K. Renner, B. Timischl, A. Mackensen, L. Kunz-Schughart, R. Andreesen, S.W. Krause, M. Kreutz, Inhibitory effect of tumor cell–derived lactic acid on human T cells, Blood. 109 (2007) 3812–3819. https://doi.org/10.1182/blood-2006-07-035972.

[30] S. Chen, X. Zhou, X. Yang, W. Li, S. Li, Z. Hu, C. Ling, R. Shi, J. Liu, G. Chen, N. Song, X. Jiang, X. Sui, Y. Gao, Dual Blockade of Lactate/GPR81 and PD-1/PD-L1 Pathways Enhances the Anti-Tumor Effects of Metformin, Biomolecules. 11 (2021) 1373. https://doi.org/10.3390/biom11091373.

[31] C.-H. Chang, J. Qiu, D. O’Sullivan, M.D. Buck, T. Noguchi, J.D. Curtis, Q. Chen, M. Gindin, M.M. Gubin, G.J.W. van der Windt, E. Tonc, R.D. Schreiber, E.J. Pearce, E.L. Pearce, Metabolic Competition in the Tumor Microenvironment Is a Driver of Cancer Progression, Cell. 162 (2015) 1229–1241. https://doi.org/10.1016/j.cell.2015.08.016.

[32] D. Chen, H.B. Barsoumian, G. Fischer, L. Yang, V. Verma, A.I. Younes, Y. Hu, F. Masropour, K. Klein, C. Vellano, J. Marszalek, M. Davies, M.A. Cortez, J. Welsh, Combination treatment with radiotherapy and a novel oxidative phosphorylation inhibitor overcomes PD-1 resistance and enhances antitumor immunity, J Immunother Cancer. 8 (2020) e000289. https://doi.org/10.1136/jitc-2019-000289.

[33] S. Ganapathy-Kanniappan, Linking tumor glycolysis and immune evasion in cancer: Emerging concepts and therapeutic opportunities, Biochimica et Biophysica Acta (BBA) - Reviews on Cancer. 1868 (2017) 212–220. https://doi.org/10.1016/j.bbcan.2017.04.002.

[34] V. Aggarwal, C.A. Montoya, V.S. Donnenberg, S. Sant, Interplay between tumor microenvironment and partial EMT as the driver of tumor progression, IScience. 24 (2021) 102113. https://doi.org/10.1016/j.isci.2021.102113.

[35] K. Skibbe, A.-K. Brethack, A. Sünderhauf, M. Ragab, A. Raschdorf, M. Hicken, H. Schlichting, J. Preira, J. Brandt, D. Castven, B. Föh, R. Pagel, J.U. Marquardt, C. Sina, S. Derer, Colorectal Cancer Progression Is Potently Reduced by a Glucose-Free, High-Protein Diet: Comparison to Anti-EGFR Therapy, Cancers (Basel). 13 (2021) 5817. https://doi.org/10.3390/cancers13225817.

[36] R. Luna-Yolba, J. Marmoiton, V. Gigo, X. Marechal, E. Boet, A. Sahal, N. Alet, I. Abramovich, E. Gottlieb, V. Visentin, M.R. Paillasse, J.-E. Sarry, Disrupting Mitochondrial Electron Transfer Chain Complex I Decreases Immune Checkpoints in Murine and Human Acute Myeloid Leukemic Cells, Cancers (Basel). 13 (2021) 3499. https://doi.org/10.3390/cancers13143499.

[37] A.M. Bolger, M. Lohse, B. Usadel, Trimmomatic: a flexible trimmer for Illumina sequence data, Bioinformatics. 30 (2014) 2114–2120. https://doi.org/10.1093/bioinformatics/btu170.

[38] A. Dobin, C.A. Davis, F. Schlesinger, J. Drenkow, C. Zaleski, S. Jha, P. Batut, M. Chaisson, T.R. Gingeras, STAR: ultrafast universal RNA-seq aligner, Bioinformatics. 29 (2013) 15– 21. https://doi.org/10.1093/bioinformatics/bts635.

[39] D. van Dijk, R. Sharma, J. Nainys, K. Yim, P. Kathail, A.J. Carr, C. Burdziak, K.R. Moon, C.L. Chaffer, D. Pattabiraman, B. Bierie, L. Mazutis, G. Wolf, S. Krishnaswamy, D. Pe’er, Recovering Gene Interactions from Single-Cell Data Using Data Diffusion, Cell. 174 (2018) 716-729.e27. https://doi.org/10.1016/j.cell.2018.05.061.

[40] L.A. Byers, L. Diao, J. Wang, P. Saintigny, L. Girard, M. Peyton, L. Shen, Y. Fan, U. Giri, P.K. Tumula, M.B. Nilsson, J. Gudikote, H. Tran, R.J.G. Cardnell, D.J. Bearss, S.L. Warner, J.M. Foulks, S.B. Kanner, V. Gandhi, N. Krett, S.T. Rosen, E.S. Kim, R.S. Herbst, G.R. Blumenschein, J.J. Lee, S.M. Lippman, K.K. Ang, G.B. Mills, W.K. Hong, J.N. Weinstein, I.I. Wistuba, K.R. Coombes, J.D. Minna, J. v. Heymach, An Epithelial–Mesenchymal Transition Gene Signature Predicts Resistance to EGFR and PI3K Inhibitors and Identifies Axl as a Therapeutic Target for Overcoming EGFR Inhibitor Resistance, Clinical Cancer Research. 19 (2013) 279–290. https://doi.org/10.1158/1078-0432.CCR-12-1558.

[41] T.Z. Tan, Q.H. Miow, Y. Miki, T. Noda, S. Mori, R.Y. Huang, J.P. Thiery, Epithelial-mesenchymal transition spectrum quantification and its efficacy in deciphering survival and drug responses of cancer patients, EMBO Mol Med. 6 (2014) 1279–1293. https://doi.org/10.15252/emmm.201404208.

[42] A. Liberzon, A. Subramanian, R. Pinchback, H. Thorvaldsdottir, P. Tamayo, J.P. Mesirov, Molecular signatures database (MSigDB) 3.0, Bioinformatics. 27 (2011) 1739–1740. https://doi.org/10.1093/bioinformatics/btr260.

[43] S. v. Puram, I. Tirosh, A.S. Parikh, A.P. Patel, K. Yizhak, S. Gillespie, C. Rodman, C.L. Luo, E.A. Mroz, K.S. Emerick, D.G. Deschler, M.A. Varvares, R. Mylvaganam, O. Rozenblatt-Rosen, J.W. Rocco, W.C. Faquin, D.T. Lin, A. Regev, B.E. Bernstein, Single-Cell Transcriptomic Analysis of Primary and Metastatic Tumor Ecosystems in Head and Neck Cancer, Cell. 171 (2017) 1611-1624.e24. https://doi.org/10.1016/j.cell.2017.10.044.

[44] L. Yu, M. Lu, D. Jia, J. Ma, E. Ben-Jacob, H. Levine, B.A. Kaipparettu, J.N. Onuchic, Modeling the Genetic Regulation of Cancer Metabolism: Interplay between Glycolysis and Oxidative Phosphorylation, Cancer Res. 77 (2017) 1564–1574. https://doi.org/10.1158/0008-5472.CAN-16-2074.

[45] D. Jia, B.B. Paudel, C.E. Hayford, K.N. Hardeman, H. Levine, J.N. Onuchic, V. Quaranta, Drug-Tolerant Idling Melanoma Cells Exhibit Theory-Predicted Metabolic Low-Low Phenotype, Front Oncol. 10 (2020). https://doi.org/10.3389/fonc.2020.01426.

[46] S. Aibar, C.B. González-Blas, T. Moerman, V.A. Huynh-Thu, H. Imrichova, G. Hulselmans, F. Rambow, J.-C. Marine, P. Geurts, J. Aerts, J. van den Oord, Z.K. Atak, J. Wouters, S. Aerts, SCENIC: single-cell regulatory network inference and clustering, Nat Methods. 14 (2017) 1083–1086. https://doi.org/10.1038/nmeth.4463.

[47] D.-B. Asante, M. Morici, G.R.K.A. Mohan, E. Acheampong, I. Spencer, W. Lin, P. van Miert, S. Gibson, A.B. Beasley, M. Ziman, L. Calapre, T.M. Meniawy, E.S. Gray, Multi-Marker Immunofluorescent Staining and PD-L1 Detection on Circulating Tumour Cells from Ovarian Cancer Patients, Cancers (Basel). 13 (2021) 6225. https://doi.org/10.3390/cancers13246225.

[48] D. Jia, M. Lu, K.H. Jung, J.H. Park, L. Yu, J.N. Onuchic, B.A. Kaipparettu, H. Levine, Elucidating cancer metabolic plasticity by coupling gene regulation with metabolic pathways, Proceedings of the National Academy of Sciences. 116 (2019) 3909–3918. https://doi.org/10.1073/pnas.1816391116.

[49] D. Jia, B.B. Paudel, C.E. Hayford, K.N. Hardeman, H. Levine, J.N. Onuchic, V. Quaranta, Drug-Tolerant Idling Melanoma Cells Exhibit Theory-Predicted Metabolic Low-Low Phenotype, Front Oncol. 10 (2020). https://doi.org/10.3389/fonc.2020.01426.

[50] F. Antonangeli, A. Natalini, M.C. Garassino, A. Sica, A. Santoni, F. di Rosa, Regulation of PD-L1 Expression by NF-κB in Cancer, Front Immunol. 11 (2020). https://doi.org/10.3389/fimmu.2020.584626.

[51] Y. Fan, T. Li, L. Xu, T. Kuang, Comprehensive Analysis of Immunoinhibitors Identifies LGALS9 and TGFBR1 as Potential Prognostic Biomarkers for Pancreatic Cancer, Comput Math Methods Med. 2020 (2020) 1–13. https://doi.org/10.1155/2020/6138039.

[52] D.P. Cook, B.C. Vanderhyden, Context specificity of the EMT transcriptional response, Nat Commun. 11 (2020) 2142. https://doi.org/10.1038/s41467-020-16066-2.

[53] K. Watanabe, N. Panchy, S. Noguchi, H. Suzuki, T. Hong, Combinatorial perturbation analysis reveals divergent regulations of mesenchymal genes during epithelial-to-mesenchymal transition, NPJ Syst Biol Appl. 5 (2019) 21. https://doi.org/10.1038/s41540-019-0097-0.

[54] M. Pillai, G. Rajaram, P. Thakur, N. Agarwal, S. Muralidharan, A. Ray, D. Barbhaya, J.A. Somarelli, M.K. Jolly, Mapping phenotypic heterogeneity in melanoma onto the epithelial-hybrid-mesenchymal axis, Front Oncol. 12 (2022). https://doi.org/10.3389/fonc.2022.913803.

[55] J.-H. Cha, W.-H. Yang, W. Xia, Y. Wei, L.-C. Chan, S.-O. Lim, C.-W. Li, T. Kim, S.-S. Chang, H.-H. Lee, J.L. Hsu, H.-L. Wang, C.-W. Kuo, W.-C. Chang, S. Hadad, C.A. Purdie, A.M. McCoy, S. Cai, Y. Tu, J.K. Litton, E.A. Mittendorf, S.L. Moulder, W.F. Symmans, A.M. Thompson, H. Piwnica-Worms, C.-H. Chen, K.-H. Khoo, M.-C. Hung, Metformin Promotes Antitumor Immunity via Endoplasmic-Reticulum-Associated Degradation of PD-L1, Mol Cell. 71 (2018) 606-620.e7. https://doi.org/10.1016/j.molcel.2018.07.030.

[56] Y. Han, D. Liu, L. Li, PD-1/PD-L1 pathway: current researches in cancer., Am J Cancer Res. 10 (2020) 727–742.

[57] X. Ju, H. Zhang, Z. Zhou, Q. Wang, Regulation of PD-L1 expression in cancer and clinical implications in immunotherapy., Am J Cancer Res. 10 (2020) 1–11.

[58] E.R. Stirling, S.M. Bronson, J.D. Mackert, K.L. Cook, P.L. Triozzi, D.R. Soto-Pantoja, Metabolic Implications of Immune Checkpoint Proteins in Cancer, Cells. 11 (2022) 179. https://doi.org/10.3390/cells11010179.

[59] L. Bornes, G. Belthier, J. van Rheenen, Epithelial-to-Mesenchymal Transition in the Light of Plasticity and Hybrid E/M States, J Clin Med. 10 (2021) 2403. https://doi.org/10.3390/jcm10112403.

[60] M.B. Morelli, C. Amantini, J.A. Rossi de Vermandois, M. Gubbiotti, A. Giannantoni, E. Mearini, F. Maggi, M. Nabissi, O. Marinelli, M. Santoni, A. Cimadamore, R. Montironi, G. Santoni, Correlation between High PD-L1 and EMT/Invasive Genes Expression and Reduced Recurrence-Free Survival in Blood-Circulating Tumor Cells from Patients with Non-Muscle-Invasive Bladder Cancer, Cancers (Basel). 13 (2021) 5989. https://doi.org/10.3390/cancers13235989.

[61] Y. Jiang, H. Zhan, Communication between EMT and PD-L1 signaling: New insights into tumor immune evasion, Cancer Lett. 468 (2020) 72–81. https://doi.org/10.1016/j.canlet.2019.10.013.

[62] S. Brabletz, H. Schuhwerk, T. Brabletz, M.P. Stemmler, Dynamic EMT: a multi-tool for tumor progression, EMBO J. 40 (2021). https://doi.org/10.15252/embj.2021108647.

[63] W. Hua, P. ten Dijke, S. Kostidis, M. Giera, M. Hornsveld, TGFβ-induced metabolic reprogramming during epithelial-to-mesenchymal transition in cancer, Cellular and Molecular Life Sciences. 77 (2020) 2103–2123. https://doi.org/10.1007/s00018-019-03398-6.

[64] F. Nakasuka, S. Tabata, T. Sakamoto, A. Hirayama, H. Ebi, T. Yamada, K. Umetsu, M. Ohishi, A. Ueno, H. Goto, M. Sugimoto, Y. Nishioka, Y. Yamada, M. Tomita, A.T. Sasaki, S. Yano, T. Soga, TGF-β-dependent reprogramming of amino acid metabolism induces epithelial–mesenchymal transition in non-small cell lung cancers, Commun Biol. 4 (2021) 782. https://doi.org/10.1038/s42003-021-02323-7.

[65] S.C. Schwager, J.A. Mosier, R.S. Padmanabhan, A. White, Q. Xing, L.A. Hapach, P. v. Taufalele, I. Ortiz, C.A. Reinhart-King, Link Between Glucose Metabolism and EMT Drives Triple Negative Breast Cancer Migratory Heterogeneity, IScience. (2022) 105190. https://doi.org/10.1016/j.isci.2022.105190.

[66] T. Shiraishi, J.E. Verdone, J. Huang, U.D. Kahlert, J.R. Hernandez, G. Torga, J.C. Zarif, T. Epstein, R. Gatenby, A. McCartney, J.H. Elisseeff, S.M. Mooney, S.S. An, K.J. Pienta, Glycolysis is the primary bioenergetic pathway for cell motility and cytoskeletal remodeling in human prostate and breast cancer cells, Oncotarget. 6 (2015) 130–143. https://doi.org/10.18632/oncotarget.2766.

[67] M. Cerezo, S. Rocchi, Cancer cell metabolic reprogramming: a keystone for the response to immunotherapy, Cell Death Dis. 11 (2020) 964. https://doi.org/10.1038/s41419-020-03175-5.

[68] M.Z. Noman, G. Desantis, B. Janji, M. Hasmim, S. Karray, P. Dessen, V. Bronte, S. Chouaib, PD-L1 is a novel direct target of HIF-1α, and its blockade under hypoxia enhanced MDSC-mediated T cell activation, ournal of Experimental Medicine. 211 (2014) 781–790. https://doi.org/10.1084/jem.20131916.

[69] C.E. Meacham, S.J. Morrison, Tumour heterogeneity and cancer cell plasticity, Nature. 501 (2013) 328–337. https://doi.org/10.1038/nature12624.

